# Dynamic Neural Encoding of Prior Expectations and Auditory Evidence Drive Perceptual Decision-Making

**DOI:** 10.1101/2025.08.11.669640

**Authors:** Sarah Tune, Iris Borschke, Jonas Obleser

## Abstract

Perceptual decision-making has been studied extensively as the integration of evidence over time. Yet how moment-to-moment fidelity of sensory encoding interacts with prior expectations in shaping our choices remains poorly understood—particularly in the auditory domain. Here we quantify how priors act at the stage of sensory encoding and how these fluctuations predict behaviour. In a human model-based EEG study, we augmented a canonical click-train evidence accumulation task with probabilistic cues to shape prior expectations and analysed all activity with trial-resolved linear encoding models. Probabilistic cues drove alpha-oscillatory EEG patterns and induced symmetric shifts in response bias but did not change temporal integration dynamics. Neural encoding strength in lateralised auditory cortex uniquely predicted single-trial choice and confidence beyond what was explained by evidence strength or cueing alone. These findings identify sensory encoding fidelity as a mechanistic, behaviourally relevant link between priors and perceptual choice, offering new understanding how neural noise and expectations jointly shape decisions under uncertainty.

## Introduction

Perceptual decisions rely on the temporal integration of sensory evidence (Gold & Shadlen, 2007). Accumulating sensory evidence over time provides an efficient strategy to mitigate perceptual uncertainty associated with individual samples of evidence (Bogacz et al., 2006; de Gardelle & Summerfield, 2011; Smith & Ratcliff, 2004). This is particularly true for the auditory domain where inherently noisy sensory signals such as human speech and environmental sounds all unfold dynamically over time.

Yet, even in relatively constrained sequential sampling tasks, human perceptual decision-making exhibits multiple sources of suboptimality. A particularly insightful example is the previously established ‘Poisson clicks’ task (Brunton et al., 2013) in which participants judge which ear receives more randomly timed auditory clicks. Variants of this paradigm have revealed systematic deviations from normative evidence integration, including temporal weighting asymmetries, choice history effects, and consistent ear biases (Gupta et al., 2024; Keung et al., 2019; Keung et al., 2020). As another potential source of suboptimality, moment-to-moment variability in the fidelity of neural sensory encoding due to fluctuating neural activity (or ‘neural noise’) have been suggested (Garrett et al., 2013; Waschke et al., 2021; Waschke et al., 2019).

Moreover, it is well established that perceptual decisions are shaped not only by sensory evidence but also by prior expectations (von Helmholtz, 1867). Seminal work in perception across modalities has shown that observers combine incoming sensory evidence with internalized beliefs and expectations about the world, effectively performing Bayesian inference to reduce uncertainty (Ernst & Banks, 2002; Stocker & Simoncelli, 2006; Weiss et al., 2002). Neuroimaging and neurophysiological studies further reveal that such expectations modulate early sensory representations and bias activity in decision-related regions to favour anticipated outcomes (Kok et al., 2013; Summerfield & de Lange, 2014; Summerfield & Koechlin, 2008).

Yet, how prior expectations impact the temporal integration of auditory information and its underlying neural encoding has never been tested in the context of this otherwise well-studied decision-making paradigm. The click trains paradigm is one of the very few genuinely auditory paradigms in the decision-making literature (Brunton et al., 2013; Hanks et al., 2015; Keung et al., 2020; Musall et al., 2023; Pagan et al., 2024), yet the neural encoding of auditory information at the heart of it has hardly been studied in satisfying detail.

With the current EEG study, we aim at closing two key gaps in the perceptual decision-making literature: First, as a novel adaptation of the Bernoulli clicks paradigm (Keung et al., 2019), we introduced a trial-by-trial probabilistic cue that altered prior expectations about the upcoming auditory input. In half of the trials, participants received an informative cue (80% valid) indicating the more likely target ear; in the remaining trials, an uninformative neutral cue was provided.

Second, in leveraging recent advances in neural encoding modelling, we explicitly focused on the analysis of trial-by-trial neural sensory encoding fidelity in auditory cortex. We sought to investigate how neural sensory encoding varies click by click in line with changing prior expectations and evidence strength, and how it links to the ensuing behavioural choices and metacognitive corollaries.

To dissect how prior expectations influence perceptual decisions, we analyse their impact across multiple stages of the perceptual hierarchy. First, psychophysical modelling allowed us to characterise the impact of informative vs. neutral cues on behaviour, focusing on changes in response bias, perceptual sensitivity, and evidence accumulation dynamics (‘temporal integration kernels’). Second, we examine the neural dynamics from which these behavioural effects emerge, specifically focusing on the neural encoding of individual click sequences in low-frequency auditory-evoked activity.

In this study, we mainly capitalize on advanced linear encoding (‘forward’) modelling of neural activity (Crosse et al., 2016; Crosse et al., 2021; Lalor et al., 2009; Wu et al., 2006). This allows us to characterize neural responses on the basis of both millisecond-to-millisecond low-level auditory evidence as well as high-level prior expectation or ongoing evidence integration processes. Additionally, we leverage an established neural marker of auditory attentional adjustments and anticipatory bias, the lateralisation of 8–12 Hz alpha power prior to and during an auditory stimulus (Kerlin et al., 2010; Müller & Weisz, 2011; Tune et al., 2021; Tune et al., 2018; Wöstmann et al., 2016).

Finally, we employ single-trial statistical modelling to explain trial-by-trial variability in perceptual decisions, and response time as a function of these neural dynamics.

Our data show that prior expectations prompt a pronounced response bias but leave temporal integration dynamics remarkably unchanged. Neurally, prior expectations induce a consistent lateralisation of pre-stimulus alpha power and act on the neural sensory encoding in lateralised auditory cortex. Most importantly, trial-to-trial variability in neural sensory encoding precision systematically and consistently shapes the ensuing perceptual choice and response time-echoed subjective confidence.

## Results

We here report on the behavioural and EEG-derived neural dynamics in a sample of healthy young individuals (*N*=32, 18–33 yrs) who performed a novel adaptation of the previously established ‘click trains’ (Brunton et al., 2013; Keung et al., 2020) auditory decision-making task. In this evidence accumulation paradigm, participants decide which ear receives more clicks in 1-sec trains of 20 clicks. Except for the first click, each click is randomly presented to either left or right ear creating trial-by-trial variability in overall evidence strength—the relative difference in right and left click counts—and its temporal evolution (Fig. 1a).

**Figure 1.**
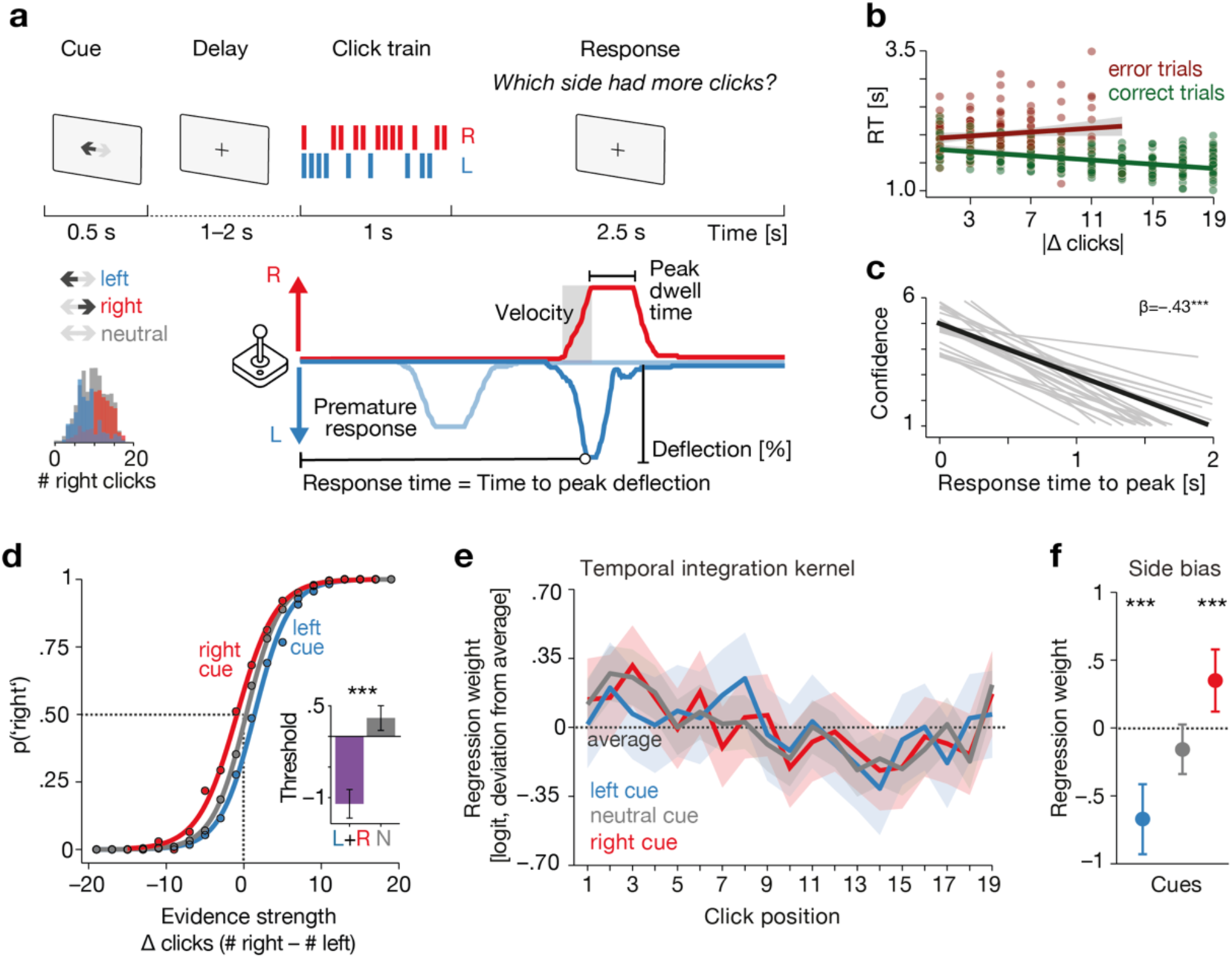
Experimental design and key behavioural results. **(a)** Trial design. Participants listened to a 1-sec streams of dichotically presented clicks and judged which side received more clicks using a joystick. Click presentation was preceded by a probabilistic cue. We derived an assay of behavioural metrics from single-trial joystick trajectories during click presentation and in the 2.5s response window. **(b)** Single-subject mean response times (coloured dots) differentially scale with increasing evidence strength for correct and incorrect trials, typically interpreted as a hallmark of confidence. **(c)** Response times significantly decrease with increasing confidence judgements in a separate behavioural replication experiment (*N*=26). Black line shows the group-level fixed effect, thin grey line subject-specific random slopes. **(d)** Informative probabilistic cues induce a symmetric shift in response bias. Inset: When modelled binarized (cued vs. neutral), cued trials shift the psychometric curve’s threshold by as much as one additional click in sensory evidence. **(e)** Temporal integration kernels fit per cue condition reveal uneven weighting of evidence over time. Regression weights from logistic mixed models predicting p(’right’) shown as deviation from the regression weight averaged across all 19 click positions along with 95% CI error bands. **(f)** Mixed model intercepts capture cue-specific side biases that align with the shift in response bias shown in panel (d). Dots indicate group-level fixed effects along with 95% CI error bars.

As a novel adaptation, we manipulated participants’ prior expectations by means of a probabilistic cue. In half of the 600 presented trials, this visual cue indicated with 80% validity which side was more likely to receive more clicks in a given trial. In the other half of trials, a neutral cue was shown. Participants reported their decision by moving a joystick toward the side they perceived to have more clicks.

From single-trial continuous joystick traces we extracted a multivariate profile of auditory decision-making, focusing on choice and response time as our key behavioural readouts. Using psychophysical analyses, we asked how variability in evidence strength and cue-driven prior expectations jointly shape auditory decision-making and its underlying temporal integration dynamics.

At the neural level, we leveraged advanced multivariate encoding models to capture trial-by-trial fluctuations in the fidelity of neural sensory representations in auditory cortex. We complemented the analysis of stimulus evoked neural responses by additionally probing how cue-induced prior expectations altered pre-stimulus neural dynamics in the 8–12 Hz alpha frequency regime.

Finally, using single-trial models of choice and response time, we aimed at explaining how moment-to-moment fluctuations in sensory encoding strength as well as pre-stimulus dynamics link to fluctuations in decision-making behaviour.

### Joystick-derived response time dynamics track both evidence and confidence

As a first step of behavioural analysis, we sought to validate our joystick-based characterization of auditory decision-making behaviour. In short, we analysed single-trial continuous joystick traces throughout click presentation and response periods to derive a set of behavioural metrics. In addition to choice, defined by the direction of maximally executed (‘peak’) joystick deflection, we extracted four additional parameters based—response time until peak deflection (termed ‘response time’ for simplicity from here on), peak dwell time, movement velocity, and deflection magnitude (Fig. 1a bottom).

Single-trial mixed-effect models revealed that all movement-related metrics except for velocity differentially scaled with the available evidence (modelled as |Δclicks|) in correct and error trials–a pattern commonly interpreted as a reflection of subjective confidence (Sanders et al., 2016). We found this confidence fingerprint to be featured most prominently in response times: in correct trials, response times decrease linearly with increasing evidence strength whereas this pattern reversed for error trials (mixed-effect model of log(response time), evidence strength × accuracy interaction: b=–0.02, standard error (SE) = 0.002, p=1.09e–33; see Fig. 1b, Fig. S1 and Table S1).

In addition, participants responded generally faster in cued compared to neutral-cued trials (left vs. neutral cue: b=–0.008, SE=0.0032, p=1.30e–2, right vs. neutral cue: b=–0.013, SE=0.0032, p=6.71e–5; see Fig. S1 and Table S1).

We further corroborated the link between our joystick-based response time measure and subjective confidence in a separate behaviour-only replication experiment (*N*=26, 20–27 years) in which participants additionally rated their confidence via button press using a 6-item scale. We found that among all joystick metrics, single-trial response time was most robustly linked to the ensuing confidence judgement (b=–2.07, SE=0.17, p=7.30e–12; note the highly consistent participant-specific random slopes in Fig. 1c, see Table S2 for full model details).

These initial results motivated our focus on response time as key metric of auditory decision-making next to single-trial choice and as a reliable proxy for confidence which we analysed next.

### Prior expectations modulate response bias but do not impact evidence accumulation dynamics

As next step in our behavioural analysis, we asked how variable evidence strength (modelled as the difference between right and left click counts, Δclicks) and probabilistic cue information impacted auditory decision-making.

First, replicating previous findings (Keung et al., 2019; Keung et al., 2020), perceptual decisions in our click trains task variant were well approximated by a logistic function of evidence strength (logistic mixed-effects model, Δclicks: odds ratio (OR)=1.63, 95% confidence interval (CI) [1.55–1.70], p=3.59e–99, see Table S3).

Second, as illustrated by the comparison of cue-specific psychometric curves in Fig. 1d, we observed that informative cues induced a symmetric shift in response bias (left vs. neutral cue: OR=0.62, CI [0.50–0.77], p=3.51e–5; right vs. neutral cue: OR=1.63, CI [1.30–2.03], p=3.51e–5). More specifically, a change in prior expectations shifted the psychometric curve horizontally by an amount equivalent to approximately one additional click of sensory evidence.

Third, we found that a cue to the right but not the left side significantly decreased perceptual sensitivity compared to the neural-cue condition (interaction sensory evidence × right vs. neutral cue: OR=0.95, CI [0.92–0.98], p=3.50e–3).

Overall, the logistic mixed-effects model accounted for approximately 80% of the variance in perceptual decisions (R^2^_conditional_=0.80).

To better understand the source of this highly systematic bias in decision-making, we then focused on the underlying temporal dynamics of evidence accumulation across cue conditions. We employed a separate set of logistic mixed-effect models to quantify and compare cue-specific temporal integration kernels. In essence, temporal integration kernels capture the relative weight each individual click position bears on the ultimate decision, with uneven kernels indicating a systematic departure from the ideal of uniform evidence weighting over time (Okazawa et al., 2018; Waskom et al., 2019).

As shown in Fig. 1e, we replicate previous findings of uneven kernel shape driven by a relative upweighting of sequence-initial clicks, followed by a gradual decline in weight, and, in our case, also an upweighting of sequence-final click information (Keung et al., 2019; Keung et al., 2020). A cubic-trend model fit the neutral-cue kernel significantly better than a flat (intercept-only) model (F(3,15)=13.683, p=1.44e–4; ΔAIC=–19.05), indicating a significant deviation from optimal, temporally unbiased evidence integration.

However, as the comparison of cue-specific kernels reveals, varying states of prior expectations leave the temporal dynamics of evidence accumulation remarkably unchanged (ORs of cue × click position ranged from 0.8–1.2, all p>0.08; see also Figure S2 for cue × position interaction kernels). In line with results from psychometric curve modelling, these logistic mixed models confirmed that participants responded unbiased in the neutral-cue condition (OR=0.86, CI [0.71– 1.03], p=9.37e–2) but exhibited a systematic side bias in the right- and left-cued condition (OR=1.45, CI [1.16–1.81], p=1.30e–3; OR=0.50, CI [0.39–0.65], p=1.18e–7).

In summary, at the behavioural level, we found that both key metrics of decision behaviour, choice and response time (as a proxy measure of subjective confidence) scale with prior expectation and available evidence.

### Prior expectations modulate lateralised pre-stimulus alpha power dynamics

We next turned to the neural level to dissect how fluctuations in prior expectations and evidence strength were reflected in neural signatures situated at distinct neural frequency regimes and temporally constrained to pre- vs. peri-stimulus periods.

We began by inspecting changes in whole-trial current source density-transformed oscillatory power within a restricted frequency range of 1–30 Hz. As shown in Fig. 2a, both visual cue and click presentation induced a pronounced 8–12 Hz alpha power desynchronization that lasted throughout the dedicated response window period. We then focused on a pre-stimulus time window (–1 to – 0.1 s relative to click train onset) to test in how far cue-driven prior expectation would lead to alpha power-mediated attentional adjustments. Given the spatial nature of our probabilistic cue and perception task, we relied on the lateralisation of alpha power as a widely established neural signature of spatial (auditory) attention (Kerlin et al., 2010; Müller & Weisz, 2011; Tune et al., 2021; Tune et al., 2018; Wöstmann et al., 2016).

**Figure 2.**
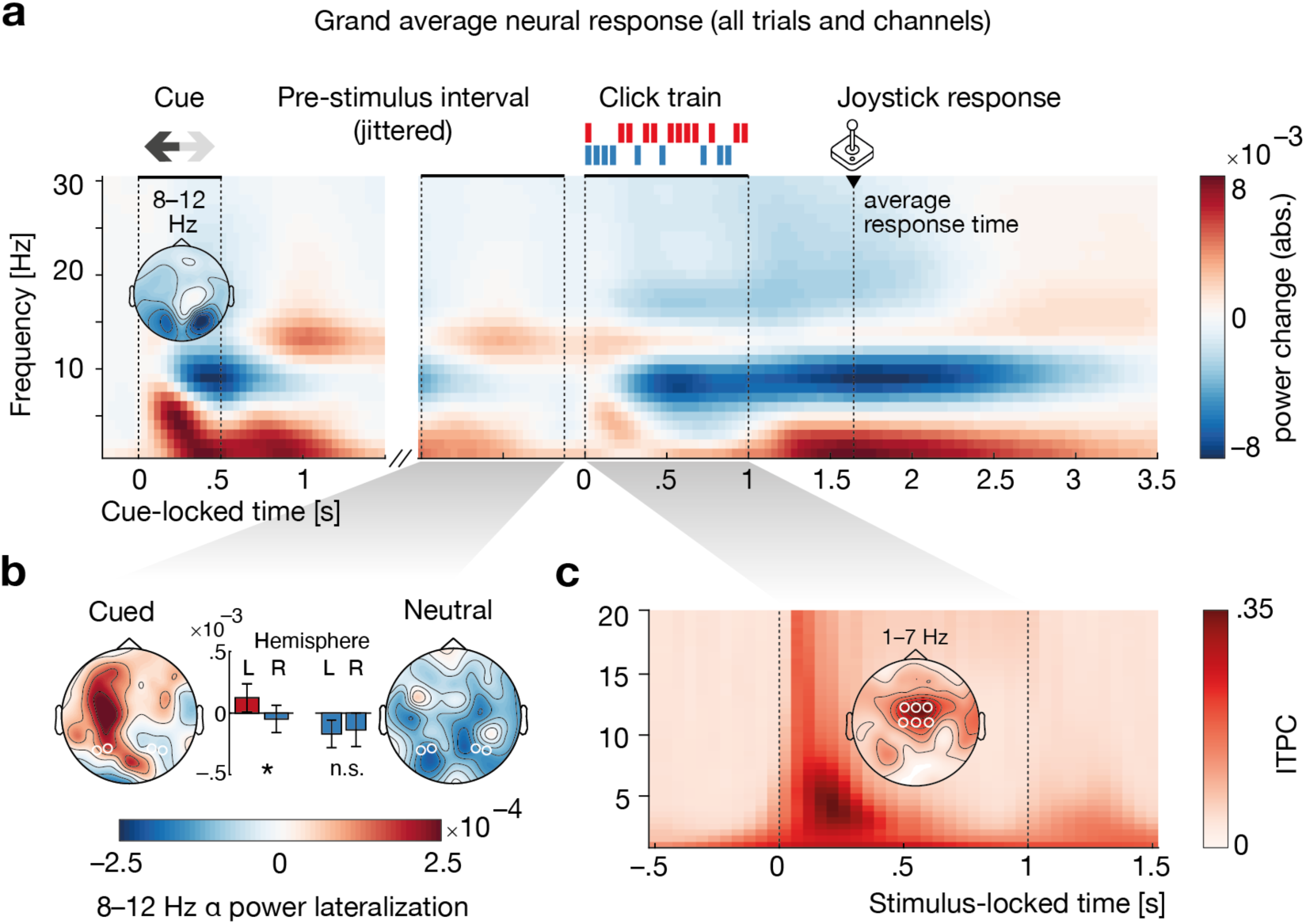
Task-related neural dynamics. **(a)** Whole-trial time–frequency representation of oscillatory power (after current source density transformation), normalized to a pre-cue baseline and averaged across all *N*=32 participants and 64 scalp channels. Topography shows 8–12 alpha power during probabilistic cue presentation. **(b)** Informative but not neutral cues induce a significant lateralisation of pre-stimulus (–1 to –0.1 s) 8–12Hz alpha power in pre-selected parietal channels (shown by white outlines). Alpha lateralisation was computed as (left – right)/(left + right), using cue direction for informative-cue trials and participants’ eventual choice for neutral-cue trials. **(c)** Inter-trial phase coherence (ITPC) averaged across all participants and six fronto-central scalp channels (shown by white outlines) in low-frequencies increases time-locked to click train onset. Topography shows 1–7 Hz ITPC during click train presentation (0–1 s).

We analysed alpha power lateralisation in pre-defined pairs of lateralised parietal channels by comparing relative power levels in cued-left–cued-right trials in the informative cue condition, and chose-left–chose-right trials in the neutral cue conditions. Indeed, we observed a weak but significant alpha power lateralisation over for informative-cue but not neutral-cue trials (Wilcoxon signed-rank test on mean hemispheric power difference, cued: z=2.00, p=0.045, neutral: z=–0.19, p=0.85, see Fig. 2b).

Click-train presentation produced a pronounced, time-locked increase in inter-trial phase coherence (1–7 Hz) over central (auditory) electrodes (Fig. 1c). This peri-stimulus ITPC surge essentially reflects the summation of auditory-evoked potentials—the stereotyped, phase-locked neural response to each individual click.

A primary aim of the current study was to characterise, on a trial-by-trial basis, how faithfully each click sequence is represented in the evoked response— and to determine how both top-down expectations and bottom-up sensory factors modulate the amplitude, timing, and spatial distribution of those phase-locked potentials. To this end, we employed advanced multivariate neural sensory encoding models reported next.

### Lateralised auditory neural responses encode sensory evidence and its modulation by prior expectations

To quantify how trial-by-trial variations in both sensory evidence and prior expectation shape auditory evoked responses, we fitted multivariate (forward) encoding models to each participant’s single-trial EEG responses filtered at 1–8 Hz. Each model treated the observed neural response as a linear sum of seven sparse ‘impulse’ regressors—click train onset and adaptation, final click offset, click side, side-change, evidence-accumulation profile, cue prior, and cue × click side interaction—each convolved with its own temporal response function (TRF). By optimizing ridge-penalized weights via leave-one-trial-out cross-validation, we obtained both the regressor-specific TRFs and a per-trial prediction accuracy index (Pearson’s r) that quantifies the fidelity with which each click sequence (and its modulation by prior expectation) is reflected in the evoked response (see Fig. 3a and Methods for modelling details).

**Figure 3.**
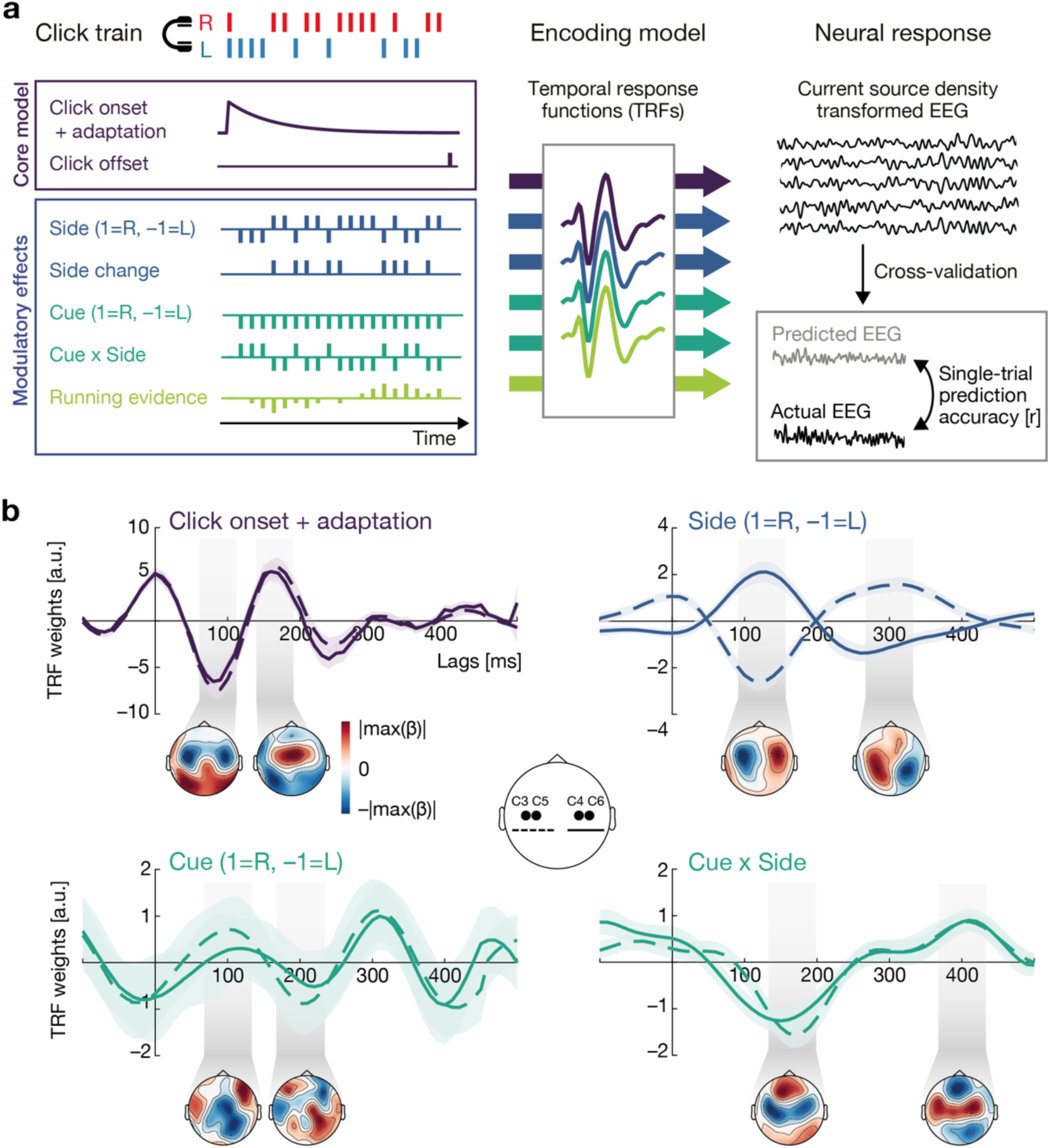
Linear forward model of neural sensory encoding in auditory cortex. **(a)** Linear encoding model details. We modelled single-trial neural sensory encoding of click sequences per EEG channel with a 7-regressor regularized linear forward model (see text and Methods for details). Using cross-validation, we estimated regressor-specific temporal response functions (TRFs) and jointly used them to predict the ensuing single-trial neural response. The correlation of predicted and actual EEG responses yielded single-trial prediction accuracy values. **(b)** Trial-average TRFs (±SEM) for four selected regressors, each averaged across all *N*=32 participants and for two pairs of lateralised central channels (dashed curves= C3, C5; solid curves= C4, C6). Note the differences in the magnitude of TRF weights across individual regressors.

As the temporal response functions of four selected regressors shown in Fig. 3b reveal, this modelling approach successfully recovered the click-onset-related canonical auditory evoked response (click onset + adaption, upper left panel) as well as its modulation by both low-level perceptual factors (click side, upper right panel) and top-down expectations (cue and cue × side, bottom panels; see Figure S3 for remaining TRFs). The click onset TRF features the overall strongest weights with prominent deflections at early time lags reminiscent of the auditory N1–P2 complex and a spatial distribution (based on current source density-transformed EEG activity) indicating, in part lateralised, auditory sources.

This canonical evoked response was modulated by click side (signed regressor: 1=right click, –1=left click): At lateralised auditory channels, clicks to the respective contralateral ear evoked generally larger early sensory responses, in particular the auditory N1, than clicks to the ipsilateral ear. This pattern reversed in direction at lags of ∼300ms, and shifted to more posterior sources in line with post-perceptual processing stages.

Perhaps unsurprisingly, the morphology and topography of the cue-driven TRF revealed a mapping between stimulus and ensuing neural response that appeared least auditory in nature, and least temporally yoked to the onset of individual clicks: instead, topographies indicate lateralised parietal to occipital sources and a quasi-rhythmic morphology suggestive of oscillatory dynamics in the alpha frequency range.

We did, however, observe that informative cues differentially affected the neural auditory encoding of individual, lateralised clicks. The cue × side interaction TRF revealed by a positive-going peak at lags ∼ 200 ms which indicates a modulation of the P2-like auditory evoked response captured by the click onset TRF: under neutral cues, lateralised clicks evoke smaller N1 and larger P2-responses in the ipsi-compared to the contralateral hemisphere. By comparison, informative cues act by specifically boosting P2 amplitude over auditory channels ipsilateral to both the cue and click presentation while leaving side-related amplitude differences in the N1 component relatively unchanged (see Fig. S4 for model-predicted single-click impulse responses).

Taken together, our forward model shows that the neural encoding of individual clicks in neural auditory activity scales with bottom-up sensory and top-down expectation-driven factors.

### Neural encoding strength reflects evidence and integration signatures

As our final and pivotal analysis step, we sought to explore whether trial-to-trial fluctuation in neural encoding strength would explain variability in perceptual choice and response time. To this end, we first analysed single-trial prediction accuracies (our proxy of neural encoding strength) derived from the full 7-regressor encoding model. As shown in Fig. 4a, computed across all trials, prediction accuracy was highest for a cluster of central auditory channels indicating a strong contribution of neural activity at these channels to the overall neural response.

**Figure 4.**
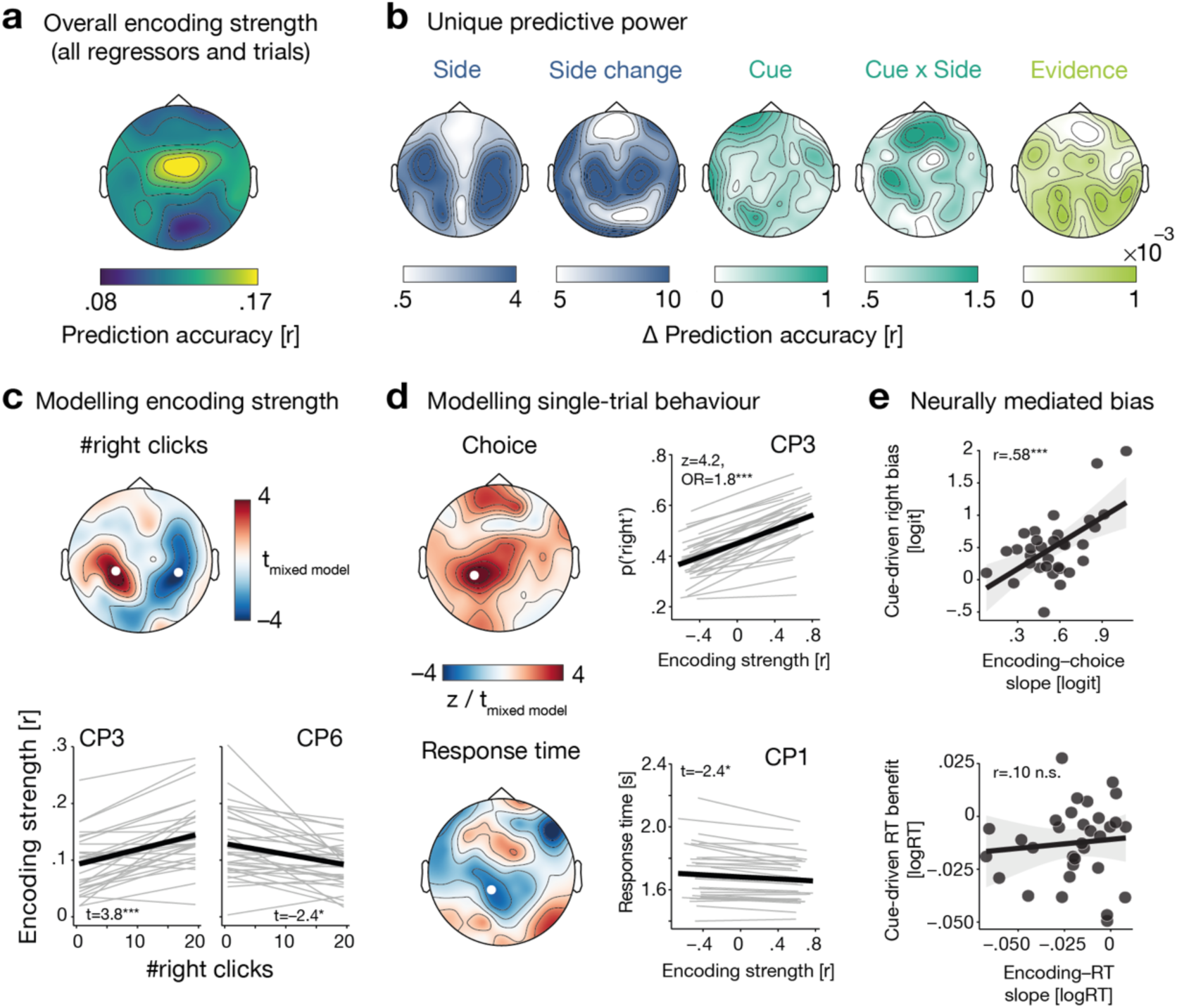
Neural encoding strength predicts single-trial behaviour. **(a)** Spatial distribution of neural sensory encoding strength (quantified via EEG prediction accuracy) averaged across all trials and *N*=32 participants. **(b)** Topographies showing spatial specificity of individual regressors’ unique predictive power. Predictive power was quantified as the respective change in prediction accuracy induced by temporally scrambling individual regressor information. **(c)** Top, spatial distribution of the group-level fixed effect of evidence strength (modelled as # of right clicks) on neural encoding strength as derived from single-trial linear mixed-modelling per channel. Bottom, Model-predicted marginal effect of evidence strength on single-trial neural sensory encoding at both the group-level (thick black line) and individual-participant level (thin grey lines showing random slopes) at the channels highlighted in white above. **(d)** Results of channel-specific linear mixed effects models linking neural encoding strength to single-trial choice and response time at the within- and between-subject level, respectively. Topographies show spatial distribution of z- and t-values associated with the within--subject effect of encoding strength after statistically accounting for cue information and evidence strength (Δclicks / |Δclicks|). Line graphs show the group-level fixed effect (thick black line) along with participant-specific random slopes (thin grey lines) for selected single channels (CP3, CP1). **(e)** Top, Strength of individual encoding–choice link is predictive of individual cue-induced right-ear side, both quantified via participant-specific random slope estimates. Bottom, Strength of encoding to response time link is statistically independent of individual cue benefit on response time.

To gain a more nuanced understanding of the unique contribution of each individual regressor to overall encoding strength, we quantified the relative reduction in prediction accuracy (Δ prediction accuracy) caused by randomly scrambling individual regressors one by one. Note that we always kept the click onset and offset regressors intact as they formed the core model capturing the canonical evoked responses.

The resultant topographies (Fig. 4b) clearly support a meaningful separation into low-level perceptual factors more closely restricted to bilateral auditory channels (side, side change, cue × side) and integrative factors such as continuous evidence accumulation processes which featured a more posterior distribution.

Using channel-specific linear mixed-effect models, we found that single-trial encoding strength significantly scaled with the number of right clicks in a given sequence (Fig. 4c): The more clicks were delivered to the right ear, the more pronounced the difference in encoding strength across bilateral central channels as higher right-click count led to better encoding over right channels and worse encoding over left channels (channel CP3: b=0.0027, SE=0.0007, p=4.75e–4; channel CP6: b=–0.0019, SE=0.008, p=1.98e–2, see also Table S4).

In sum, these distinct, spatially and temporally dissociable feature contributions underscore that our multivariate encoding framework does more than recover generic evoked responses—it meaningfully parses the rich, click-by-click dynamics of perceptual encoding and evidence integration that would be obscured by traditional ERP analyses.

### Neural encoding dynamics mediate cue-induced response bias

Having ensured the meaningfulness of our derived single-trial neural encoding measure, we turned to the final step of our analysis that linked fluctuations in encoding strength to fluctuations in decision-making behaviour. More specifically, we asked to which degree single-trial states of neural encoding would explain the behavioural outcome over and above the known effects of cue-driven prior expectations and given evidence strength.

To this end, we fit channel-wise (generalized) linear mixed-effect model of choice and response time that included both within- and between-subject regressors of encoding strength (see Methods for details) as well as cue information, evidence strength and their interaction.

We observed that both key behavioural readouts, choice and response time, were meaningfully and robustly linked to within-subject states of neural encoding fidelity: Trials with relatively higher encoding precision over left central-parietal channels significantly increased the probability of reporting more clicks on the right ear (channel CP3: OR=1.75, CI [1.35–2.28], p=8.13e–5, Fig. 4d, top channel) while also decreasing response time (channel CP1: b=–.02, SE=0.008, p=3.51e–2; Fig. 4d, bottom panel, and Table S6). Notably, in terms of effect size, the odds ratio for this neural predictor was on par with those observed for the two informative cue conditions (see Table S5 for full model details).

Moreover, both brain-behaviour relationships were highly consistent across individual participants and were restricted to the within-subject state-level, which offers the most compelling evidence for any brain-behaviour link. (See Fig. S5 for absence of between-subject, trait-levels effects.)

Following up on these results, we asked in how far inter-individual differences in the strength of these observed brain–behaviour relationships and in cue-induced modulations of behaviour were connected. Put differently, we probed the extent to which cue-induced response bias shifts or response time decreases were neurally mediated by encoding dynamics. To answer this question, we estimated participant-specific random slopes of within-subject encoding strength and right vs. neutral cue and correlated them per behavioural metric (Fig. 4e). The strength of the encoding–behaviour relationship explained inter-individual differences in right cue-induced response bias (r=0.58, p= 5.09e–4) but not response time benefit (r=0.09, p=0.62), indicating that the rightward shift in response bias is at least in part mediated by cue-informed neural encoding fidelity.

Finally, we asked whether the pre-stimulus neural state, quantified by the degree of single-trial parietal alpha power lateralisation (ALI) would predict behaviour; independently or in interaction with the respective state of peri-stimulus neural encoding. When modelling choice, we observed a trend toward an interactive effect of pre- and peri-stimulus dynamics: relatively high level of alpha power lateralisation further boosted the positive relationship of encoding strength on the probability of perceiving more clicks to the right ear (OR=1.25, CI [0.99–1.58], p=1.03e-1). In addition, participants who showed a generally more pronounced alpha lateralisation also responded faster (b=–0.003, SE=0.001, p=2.54e–2). Note, however, that state-level changes in pre-stimulus alpha lateralisation and neural encoding strength were themselves statistically unrelated (channel CP3: b=–0.0018, SE=0.0015, p=2.20e–1; channel CP1: b=–0.0008, SE=0.0015, p=5.63e–1).

In sum, our results show that the millisecond-by-millisecond fidelity with which the brain encodes each click sequence carries unique, behaviourally meaningful information—over and above the effects of cueing and evidence strength—about both the eventual choice and a response time-encoded proxy of subjective confidence.

## Discussion

How the auditory system integrates sensory information under uncertainty—and how this process is shaped by prior expectations—remains incompletely understood, especially at the level of early sensory encoding. Here, we combined a canonical click-train evidence-accumulation task with trial-by-trial probabilistic cues and multivariate EEG encoding models to quantify, with millisecond precision, how priors act on the neural representation of auditory evidence.

Behaviourally, informative cues shifted response bias by the equivalent of one additional click of evidence while leaving the temporal profile of evidence accumulation unchanged. Neurally, priors induced lateralisation of pre-stimulus alpha power and modulated the fidelity with which individual clicks were encoded in lateralised auditory cortex.

Crucially, trial-resolved fluctuations in this encoding fidelity predicted both perceptual choice and a response-time proxy of subjective confidence, linking prior expectations to behavioural bias via changes in sensory encoding rather than altered accumulation dynamics. We will now discuss these findings in more detail.

### Dissociable modulation of response bias and accumulation dynamics

Perceptual decisions in our task revealed two distinct signatures of suboptimality: informative cues produced shifts in response bias. Also, we replicate the “uneven” psychophysical kernels reported by Keung et al. (2019, 2020). Crucially, however, this shape of the accumulation kernel was largely immune to cueing.

This dissociation suggests that overall decision bias and fine-grained integration dynamics are mediated by separable mechanisms. Indeed, a recent biophysically-plausible cortical circuit model of decision-making, based on dynamic divisive normalization as a canonical neural coding principle (Carandini & Heeger, 2011), accommodates this separation by fitting independent parameters for response bias versus kernel shape (Keung et al., 2020). More generally, cortical circuit modelling suggests that the shape and dynamics of psychophysical kernels are consistently and mechanistically linked to the interplay of cortical excitatory and inhibitory forces (Cheadle et al., 2014; Lam et al., 2022; Murphy et al., 2021; Okazawa & Kiani, 2023; Okazawa et al., 2018; Wimmer et al., 2015).

By contrast, prior knowledge about upcoming stimuli and thereby in our task also the probable best choice may exert their influence via more global modulatory signals. These signals could shift the decision criterion or apply a uniform gain in sensory processing without altering click-position–specific integration dynamics. In canonical models of perceptual decision-making such as the drift–diffusion model (DDM), such effects are often formalised as pre-stimulus criterion shifts (starting-point bias) or constant changes in drift rate that leave the relative temporal weighting of evidence unaffected (Gold & Shadlen, 2007; Ratcliff & McKoon, 2008; Ratcliff & Smith, 2004).

Neurophysiological work demonstrates that such biases can manifest well before stimulus onset, for example in the form of elevated preparatory activity in motor and decision-related regions when a response is expected (de Lange et al., 2010; de Lange et al., 2013; Donner et al., 2009; Gould et al., 2012). Other studies combining behavioural modelling with neural data suggest that prior expectations can also accelerate the accumulation of sensory evidence in favour of the predicted alternative (Forstmann et al., 2010; Mulder et al., 2012; St John-Saaltink et al., 2016), possibly reflecting top-down modulation of early sensory representations (Kok et al., 2013; Wyart, Nobre, et al., 2012).

This converging evidence motivates a critical question: to what extent do expectation-driven biases originate during the encoding of sensory evidence itself, before the integration process begins? Our neural-encoding–model results provide much-needed neurophysiological evidence from the comparatively understudied auditory domain, capturing how prior information shapes the encoding of sensory inputs.

### Perceptual evidence and prior expectations converge in auditory encoding dynamics

Our forward-modelling approach exploited a key strength of multivariate TRF analysis: the ability to separate overlapping neural responses into temporally and functionally distinct components. By explicitly incorporating auditory-specific features such as adaptation to repeated clicks (Näätänen & Picton, 1987; Plomp, 1964) and heightened sensitivity to perceptual updates (Näätänen et al., 2007) alongside top-down regressors, the model captured how prior expectations act on lateralised auditory encoding at millisecond resolution.

While well-suited to isolating stimulus-evoked auditory cortical responses, it was not optimised to capture the neural representation of high-level information such as cue information itself or evidence accumulation profiles. These TRFs and their unique predictive contributions showed clearly non-auditory occipital–parietal patterns consistent with spatial auditory attention and evidence integration dynamics (Erb & Obleser, 2020; Strauß et al., 2014; Tune et al., 2018; Wöstmann et al., 2016; Wyart, de Gardelle, et al., 2012).

Our results reinforce the view that sensory encoding is not purely feed-forward: prior knowledge can shape early cortical processing of incoming evidence (Egner et al., 2010; Kok et al., 2013; Obleser, 2025; Rahnev et al., 2011; Summerfield & de Lange, 2014; Summerfield & Koechlin, 2008; Walsh et al., 2024; Wyart, Nobre, et al., 2012).

Crucially, the interaction of cue information with lateralised clicks revealed a selective modulation of the P2-like component, amplifying responses to unexpected stimuli and attenuating responses to expected stimuli, while leaving the earlier, obligatory N1 largely unchanged. This dissociation suggests that expectations do not alter the initial, stimulus-driven registration of sounds but bias more malleable later stages of selection and suppression (Crowley & Colrain, 2004; Picton, 2013; Rockstroh et al., 1992; Ross & Tremblay, 2009; Tremblay et al., 2014). Increased inhibition via amplified P2-responses enhance processing differences between expected and unexpected stimuli, and potentially gate unexpected inputs from entering downstream decision circuits (Foxe & Snyder, 2011; Luck et al., 1996; Schroger et al., 2015; Woldorff et al., 1987).

By demonstrating such component-specific modulation in the auditory domain, our results provide neurophysiological evidence that probabilistic cues can influence the sensory evidence stream before it reaches the accumulation stage. Most importantly, our study provides a mechanistic link between perceptual priors, sensory encoding precision, and biased decision-making. These data show how trial-by-trial variability in neural encoding fidelity mediates biased perceptual choices and links to response time-encoded subjective confidence.

### Neural encoding dynamics predict biased perceptual decisions

Our data not only revealed that auditory encoding is shaped by both sensory evidence and prior expectations. Also, trial-by-trial variability in encoding precision was meaningfully connected to ensuing decisions and experienced uncertainty (Krakauer et al., 2017; Tune & Obleser, 2024; Waschke et al., 2021). This relationship emerged at the state level, that is, within participants across trials, representing the strongest form of evidence for a causal, mechanistic link from neural variability to the behavioural outcome (Pernet et al., 2011; Tune et al., 2021).

Specifically, stronger encoding in the left hemisphere was associated with an increased probability of reporting more clicks on the right ear (Fig. 4d). Strikingly, individuals who exhibited a larger cue-driven shift in response bias toward the right ear also showed a stronger brain–behaviour coupling in this channel, indicating that the observed, cue-induced bias in decision-making was in part mediated by trial-by-trial fluctuations in auditory encoding precision.

These findings support an account of neural variability as not mere background noise, but as a key driver of behavioural variability in perceptual decisions and their metacognitive evaluation (Garrett et al., 2011; Garrett et al., 2013; Iemi et al., 2017; Iemi et al., 2021; Samaha et al., 2020; Waschke et al., 2021; Waschke et al., 2019; Waschke et al., 2017; Wöstmann et al., 2019).

The present results speak to a broader question in perceptual neuroscience: what is the functional role of neural variability and the associated precision of sensory representations for perceptual decision-making and its metacognitive corollaries (Obleser, 2025)?

Dissociable from slow neural oscillations (Henry & Obleser, 2012; Spitzer et al., 2017; Wyart, de Gardelle, et al., 2012), changes in cortical excitation–inhibition (E:I) balance have been linked to variable neural encoding (Cheadle et al., 2014), selective behavioural impairments (Lam et al., 2022), and to distinct sources of decision uncertainty, including sensory, inference, and selection noise (Salvador et al., 2022). Perturbations of E:I balance through pharmacological challenges (Cortes-Briones et al., 2025; Kunze et al., 2025; Salvador et al., 2022; Tune et al., 2025) provide a powerful approach to probing these links: modelling and empirical work suggests that shifts toward elevated excitation bias decisions toward impulsivity, whereas increased inhibition promotes hesitancy and indecision (Lam et al., 2022). Our adaptation of the click trains task and the proposed modelling framework are well-suited to such interventions. Beyond pharmacology, the paradigm offers a tractable model to investigate the neural underpinnings of pathological perceptual biases (Murray & Wang, 2018), such as auditory hallucinations in schizophrenia, where aberrant weighting of sensory evidence and priors is thought to arise from altered E:I balance and heightened internal noise (Adams et al., 2016; Adams et al., 2018; Buxbaum et al., 2007; Erb et al., 2020; Jardri & Deneve, 2013; Powers et al., 2017).

### Is the auditory system particularly prone to right-side biases?

The above results raise the question of why cue-induced biases were especially pronounced for right-ear choices. Previous work has suggested that, in the absence of explicit cues, the auditory system can exhibit a subtle intrinsic preference for right-ear input—likely reflecting left-hemisphere specialisation for processing rapid temporal features, even in non-speech sounds (Brown & Nicholls, 1997; Payne et al., 2016; Tanaka et al., 2021). This right-ear advantage is well-documented for dichotic listening to speech (Alavash et al., 2019; Tune et al., 2021; Tune et al., 2018). While weaker and less consistent for non-verbal stimuli, it nonetheless suggests an asymmetric operating point of the auditory system. In our neutral-cue condition, we found no such baseline bias, possibly because the presence of prior information across the experiment reset or even over-corrected this intrinsic tendency (Keung et al., 2019).

However, when cues explicitly favoured the right ear, the shift in response bias was accompanied by a reduction in perceptual sensitivity, a pattern consistent with a re-emergence of a right-ear bias. At the neural level, two mechanisms appeared to support this shift: (i) generally stronger sensory encoding of right-ear clicks in the left hemisphere, and (ii) a stronger within-subject coupling between encoding strength at these channels and the probability of reporting more clicks to the right. Together, these results suggest that both asymmetries in baseline auditory encoding and the modulatory effects of prior expectations can converge to push the system toward its default, right-ear–favouring state.

In line with this idea, we also observed a consistent pre-stimulus alpha lateralisation when informative cues were presented, but not under neutral cueing (Fig. 2b). Given the established role of 8–12 Hz alpha power in gating auditory input via spatial attention (Kerlin et al., 2010; Wöstmann et al., 2016), this suggests that anticipatory attentional biases may have further amplified the already privileged status of right-ear input. Notably, alpha lateralisation predicted faster responses across participants, indicating its functional relevance. Together with asymmetric evoked encoding and stronger encoding–choice coupling, these pre-stimulus dynamics provide converging neural evidence for a right-hemisphere attentional “set” that interacts with the auditory system’s baseline asymmetry to favour right-ear reports under predictive conditions.

## Conclusion

The fidelity of neural encoding in auditory cortex forms a dynamic bridge between prior expectations and perceptual choice. By linking trial-resolved encoding fidelity to both choice and confidence, we identify a mechanistic pathway through which prior expectations act on perception. This framework extends the decision-making literature by positioning sensory encoding fidelity—rather than only accumulation dynamics—as a key and potentially malleable determinant of behaviour under uncertainty.

## Materials and Methods

### Participants and procedure

*N*=32 healthy, right-handed individuals aged 18 to 33 years (median age = 24 yrs; 24 female) took part in a single experimental session lasting approximately 3.5 hours. All individuals reported normal or corrected-to-normal vision and no history of any neurological or psychiatric disorders. All individuals had normal hearing as assessed by pure-tone audiometric testing (Equinox 2.0, Interacoustics; mean pure-tone average of 0.5, 1, 2, and 4kHz across both ears was 5.6 ± sd 3.8 dB).

Following audiometric testing, participants performed ten blocks of an auditory decision-making task during which their electroencephalogram (EEG), electrocardiogram (ECG), blood pulse, skin conductance, and respiratory activity were recorded.

Data from five additional individuals were excluded from all analyses due to either technical problems in EEG recording (*N*=2), incomplete data acquisition (*N*=1), or because the joystick-based analysis of behavioural responses yielded less than 50 % valid trials (*N*=2).

Participants gave written informed consent and received financial compensation (10€ per hour) or course credit. Procedures were approved by the ethics committee of the University of Lübeck and were in accordance with the Declaration of Helsinki.

### Bernoulli click trains task

We used an adaptation of a previously established auditory decision-making paradigm (Brunton et al., 2013; Keung et al., 2019; Keung et al., 2020). On each trial, participants heard a 1-second train of 20 clicks through in-ear headphones (3M EARTONE Gold 3A). The first click was played to both ears, while each subsequent click was assigned to either the left or right ear according to a Bernoulli process. Participants decided which side received more clicks. They responded by moving a joystick positioned on a table in front of them (Extreme 3D Pro Precision, Logitech International S.A.) in the respective direction.

Each click was constructed as a sum of pure tones (2, 4, 6, 8, and 16Hz) convolved with a cosine envelope 3 ms in width and presented at 48 kHz. Stimulus presentation was carried out with Psychtoolbox in Matlab 2021b.

As our key adaptation, we manipulated prior expectations via a preceding probabilistic visual cue indicating which side was more likely to receive more clicks in a given trial. In 50 % of trials, an informative cue indicated that more clicks were likely played to the left or right side, respectively. In the other 50 % of trials, a neutral cue was shown, indicating that is was unknown which side would receive more clicks. Informative cues were 80 % valid but participants were not explicitly informed of this.

For every trial, we randomly selected one of five probabilities of receiving a click to the right ear (p_right_ = 0.3, 0.4, 0.5, 0.6, or 0.7), and used this probability to stochastically determine the side of each remaining click. Following this procedure, approximately 93 % of trials presented between 4 to 15 trials to a given side with only rare occurrences of more extreme asymmetry in click count between the two sides (see Fig. 1A for cue-specific distributions).

To implement our cue manipulation, per participant, we constructed all *N*=600 sequences at the beginning of the experiment. For 80 % cue validity in cued trials, we then randomly picked 120 trials with more clicks to the respective cued side, along with 30 trials with more clicks to other side. The order of spatially-cued and neutral trials in a given block was presented randomly whereas the proportion of neutral (50 %) and cued trials (25 % per side) stayed constant. Please note that due to the inadvertent use of Matlab’s default random-number seed on startup, 26 of the 32 participants received the same trial sequence.

The probabilistic visual cue consisted of a grey double-headed arrow shown centrally against a white background for 500 ms. For the left and right cue variants, the arrow was symmetrically divided in two shades of grey with the darker shade indicating the side that was more likely to receive more clicks.

Visual-cue presentation was followed by a pre-stimulus jitter period [1–2s, mean:1.5s] showing a black central fixation cross. The fixation cross remained on screen throughout the subsequent during auditory presentation and the immediately following response window of 2.5 sec. Single trials were separated by a jittered inter-trial interval [1–2.5s, mean=1.75s]. No feedback was given.

Following a short training block of 20 trials, participants performed a total of 600 trials, divided into 10 blocks of 60 trials with self-paced breaks in between.

### EEG and physiological recordings

We recorded participants’ EEG from 64 active electrode mounted to an elastic cap (ActiCap/ActiChampPlus; Brain Products, Gilching, Germany). The EEG was recorded at a sampling rate of 1000 Hz, and referenced on-line to the left mastoid electrode (TP9, ground: AFz). Impedances were kept below 20 kΩ.

As additional peripherical physiological measures, we recorded individuals’ electrocardiogram (ECG), blood pulse, skin conductance, and respiratory activity. ECG was recorded using a 3-lead configuration with one electrode were placed underneath each clavicle bone and a third electrode on the participant’s lower left abdomen. A blood pulse sensor was placed on the left index finger, while skin conductance was measured from two galvanic skin response sensors placed on the left middle and ring finger, respectively. Finally, respiratory activity was measured by means of a respiratory chest belt worn just below the sternum. All peripherical measurements were recorded as auxiliary channels along with the EEG.

### Joystick recording and analysis

We recorded single-trial joystick movements along the x-, and y-axis for the duration of the entire trial. The joystick’s neutral position was calibrated at the beginning of the experiment. Movement in either direction was converted to proportions of the maximal deflection possible, resulting in values bounded between [–1,1], accordingly.

In our analysis of behavioural responses from single-trial joystick traces we primarily focused on a 3.5s time window starting with click train presentation until the end of the ensuing 2.5 s response window. This choice was motivated by initial analyses revealing that in ∼10% of trials, participants initiated joystick movements already during click train presentation. As these ‘early response’ trials may be particularly informative for the analysis of gradual evidence accumulation and decision-making, we decided to include the auditory stimulation period in our joystick trace analyses.

We analysed participants’ behavioural decisions based on the direction and magnitude of joystick deflection. Valid responses were defined by a joystick deflection of at least 20% in either direction paired with a response time of at least 800ms post auditory stimulus onset. The latter criterion was motivated by the assumption that individuals needed to perceive at least half of the click train sequence (i.e. the first 500 ms) to gather enough evidence for a decision, plus another ∼300 ms to initiate joystick movement. This procedure resulted in the exclusion of 647 (3.4 %) timeout trials across participants.

As our primary behavioural metric of interest, we derived single-trial binary choice from the direction of maximally executed (‘peak’) joystick deflection. Note, however, that this definition of choice is highly correlated with an alternative procedure that analyses the direction in which the 20 % deflection point is first surpassed (Szul et al., 2020)(r =.98, p <.001). We also analysed response times until peak deflection, the magnitude of deflection, and how long the joystick was held at the maximally executed deflection (peak dwell time). Finally, we calculated velocity of joystick movement as executed deflection per time taken.

### Statistical analysis

We performed group-level psychophysical modelling of single-trial perceptual decisions by estimating logistic linear mixed-effects models with the package *lme4* of the statistical analysis software R. We modelled the probability of choosing ‘right’ as a function of single-trial sensory evidence strength ($clicks, # right clicks – # left clicks), prior information (neutral, left, right cue), and their interaction. In such a model, the regression weight for the sensory evidence regressors estimates the slope of the psychometric curve, that is, perceptual sensitivity. The weight of the cue regressor estimates horizontal shifts of the psychometric curve, reflecting prior-induced changes in decision bias. Lastly, the interaction of sensory evidence and prior information estimates cue-related modulation of perceptual sensitivity (Waschke et al., 2019). To account for inter-individual variability, we included participant-specific random intercepts and random slopes of sensory evidence and prior information.

Prior to fitting group-level models, we performed goodness-of-fit checks by estimating individual psychometric curves for each participant’s data using the *quickpsy* package (Linares & López-Moliner, 2016). Specifically, using only neutral-cued trials, we modelled the probability of choosing ‘right’ as a logistic function of $clicks using 1000 bootstrap iterations to estimate parameter confidence intervals and residual deviance. The observed deviance was compared against the bootstrap distribution to test whether the fitted logistic model adequately captured the data. In all cases, the residual deviance was non-significant (p>.13), indicating that the logistic function provided a reasonable fit for each participant’s neutral-cue data.

In auxiliary analyses, we used (generalized) linear mixed-effect models to investigate how response time, velocity, deflection magnitude, and peak dwell time changed in line with provided sensory evidence (here modelled as |$clicks|) and prior information. These models also included for accuracy as well as its interaction with the available evidence strength.

Brain-behaviour models were performed per individual scalp channel. As predictor they included as before the available evidence strength ($clicks for choice; |$clicks| for response times), cue information, along with neural variables split into a between- and within-subject regressor. The between-subject regressor quantifies the individual, trait-level mean neural encoding and alpha lateralisation strength across all trials, whereas the within-subject regressor quantifies the single-trial, state-level deviations from this mean (Bell et al., 2018; Tune et al., 2021; Tune & Obleser, 2024).

Across all analyses, we used simple effect coding for the categorical cue regressor (with the neutral cue as reference level), and mean-centred all continuous regressors. P-values for individual model terms in general linear mixed-effect models are based on the Satterthwaite approximation for degrees of freedom, and on z-values and asymptotic Wald tests for the generalised linear mixed-effect model of accuracy (Luke, 2017).

### Temporal integration kernel

We used another set of logistic linear mixed-effect model to quantify the temporal dynamics of evidence accumulation, and how they may be modulated by changes in cue-induced prior expectation. Specifically, per cue condition, we modelled the temporal integration kernel that characterizes the impact of each individual click position on the perceptual decision. A flat kernel indicates that all click positions have equal weight on the decision, any deviation in shape point towards a biased weighting of evidence over time (Keung et al., 2019; Keung et al., 2020).

For each of the 19 click positions following the initial click played to both ears, we coded whether the respective click was played to the right ear (right = 0.5, left = –0.5). Deviation from a flat integration kernel was assessed by expressing individual weight as deviation from the average across all 19 click positions. The intercept of such a model quantifies the position-independent probability of choosing right thereby providing an estimate of overall side bias. We included participant-specific random intercepts to account for inter-individual variability in side bias. We refrained from analysing any sequential (‘choice history’) effects, as our trial-level cue manipulation may have interfered with any tendency to base current decision-making on past choices.

Lastly, to more directly test whether prior cue information modulated evidence accumulation over time, we ran an additional model that for each click position included the interaction of cue information and direction of each click.

### Electroencephalography (EEG) analysis

Continuous EEG data were pre-processed using EEGlab and Fieldtrip toolboxes, together with customized Matlab scripts. Independent component analysis (ICA) using EEGlab’s default runica algorithm was used to identify all non-brain signal components including eye blinks and lateral eye movements, muscle activity, heartbeats and single-channel noise. Prior to ICA, EEG data were re-referenced to the average of all EEG channels (average reference), down sampled to 300 Hz, filtered between 1 and 100 Hz, and cut into 1 s epochs. Non-brain signal components were automatically identified based on EEGlab’s pre-trained IClabel algorithm (Pion-Tonachini et al., 2019); probability threshold per signal category set to p = .9) and subsequently removed from the continuous raw EEG signal.

For the analysis of neural sensory encoding, the cleaned continuous EEG data were down-sampled to f_s_=100 Hz and filtered between f_c_=1 and 8 Hz (FIR filters, zero-phase lag, order: 8f_s_/f_c_ and 2f_s_/f_c_, Hamming window). We transformed voltage data to current source density using individual digitized electrode positions and spherical splines interpolation (Kayser & Tenke, 2015). We then extracted single-trial epochs of –0.5 to 1.5 s relative to the onset of the first click as input to linear encoding modelling.

To visualize whole-trial changes in neural responses and to analyse pre-stimulus alpha power lateralisation, cleaned continuous EEG data were high-pass-filtered at 0.3 Hz (finite impulse response (FIR) filter, zero-phase lag, order 5574, Hann window) and low-pass-filtered at 180 Hz (FIR filter, zero-phase lag, order 100, Hamming window). The EEG was cut into epochs of –1 to 2 s relative to the onset of the probabilistic cue, as well as epochs of –2 to 4 s relative to click train onset to focus on fluctuation in pre- and peri-stimulus neural responses, respectively. Data were down-sampled to f_s_=250 Hz, and the same current source density transformation was applied. We computed time–frequency representations using sliding-window multi-taper convolution. We applied a Hanning taper, with a fixed 500 ms window at each frequency (1–30 Hz), advancing in 40 ms steps.

Absolute power was calculated as the square-amplitude of the spectro-temporal estimates. Since oscillatory power values typically follow a highly skewed, non-normal distribution, we applied a nonlinear transformation of the Box-Cox family (power*_trans_* = (power^p^ −1)/p with p=0.22) to minimize skewness and to satisfy the assumption of normality for parametric statistical tests involving oscillatory power values. To illustrate whole-trial neural dynamics, we then computed changes in power relative a pre-cue (–0.8–0.2 s) baseline period.

Inter-trial phase coherence (ITPC) was computed via wavelet convolution for frequencies from 1 to 20 Hz sampled in 0.5 Hz steps, and time-points from –1 s to +2 s relative to stimulus onset in 50 ms increments. ITPC at each channel– frequency–time bin was computed by normalizing each trial’s complex spectrum to unit length, summing across trials, and dividing the resultant vector magnitude by the number of trials.

### Single-trial sensory encoding models

Prior to the estimation of temporal response functions, we z-scored the EEG data channel-wise across all trials to remove inter-channel gain differences while preserving trial-to-trial fluctuations. We constructed a sparse stimulus design matrix for each trial, consisting of seven regressors sampled at 100 Hz.

The first regressor modelled sensory adaptation as a smooth exponential decay (τ = 200 ms, REFS). The second regressor captured any stimulus offset responses impulse 50 ms after the final click. The third and fourth regressors encoded click-side (+1 for right vs. –1 for left side, first click coded as zero) and cue condition (+1 for right- vs. –1 for left-cued trials, zero for neutral), respectively, coded as impulses at each click onset. The fifth regressor formed their point-wise interaction (cue×side). The sixth regressor represented signed evidence accumulation (‘running difference’) computed as the cumulative sum of side codes up to each click and then z-scored across all trials. Finally, the seventh regressor marked perceptual changes with impulses of magnitude 1 at any click where the side switched.

Linear encoding models were fit using the mTRF Toolbox, (Crosse et al., 2016) in Matlab in forward mode over time lags from –200–500 ms, and subsequently tested using more restricted lags between 0–400ms. For each subject, ridge regularization λ was selected from a grid of 10⁻⁶ to 10⁶ using cross-validation, optimizing the mean Pearson correlation between predicted and held-out EEG in an auditory fronto-central auditory region of interest (FC5, FC3, FC1, FCz, FC2, FC4, FC6, C5, C3, C1, Cz, C2, C4, C6). The chosen λ was then used to train a final encoding model on all trials. Single-trial prediction accuracy was quantified as the Pearson r between predicted and observed EEG for each of 64 scalp channels using leave-one-out cross-validation. This analysis pipeline yielded spatio-temporal TRF weights for each regressor and trial-level metrics of encoding strength for subsequent statistical analysis.

To determine the unique predictive power of individual regressors, using 5-fold cross-validation, we quantified the reduction in prediction accuracy after temporally scrambling one regressor at a time, thus holding model complexity constant but removing regressor-specific information (Musall et al., 2019).

### Attentional modulation of alpha power

To quantify cue-related changes in pre-stimulus 8–12 Hz alpha power, we calculated the single-trial, temporally resolved alpha lateralisation index as follows(Tune et al., 2021):

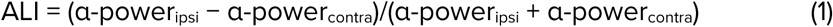

Note that for informative-cued trial, the formula included power values in the hemisphere ipsi- and contralateral to the given cue direction, whereas for neutral-cued trials, it included power values in the hemisphere ipsi- and contralateral to the eventual choice given.

To account for overall hemispheric power differences that were independent of cue effects, we first normalized single-trial power by calculating per channel, frequency and time point the power averaged across all trials and subtracted it from single trials. We then used a robust variant of the index that applies the inverse logit transform [(1 / (1 + exp(–x))] to both inputs to scale them into a common, positive-only [0;1]-bound space prior to index calculation.

For statistical analysis of cue-driven alpha lateralisation, we then averaged the alpha lateralisation index values within pre-specified lateralised parietal electrode sections (P3 & P5 vs. PC4 & P6) and extracted single-trial mean values within a pre-stimulus window of –1 to –0.1 s prior to click train onset.

## Data and code availability

The complete dataset associated with this work including raw data, EEG data analysis results, as well as corresponding code will be publicly available at OSF (https://osf.io/6y9su/).

## Acknowledgments

ST and JO are funded by the University of Lübeck. Research was supported through the German Research Foundation (DFG; TU 772/2-1; Ob 352/2-2). Nikolai Dürrbeck and Mirjam Kossmann helped implement the paradigm and acquire the data.

## Supplemental information for

### Supplemental figures

**Figure S1:**
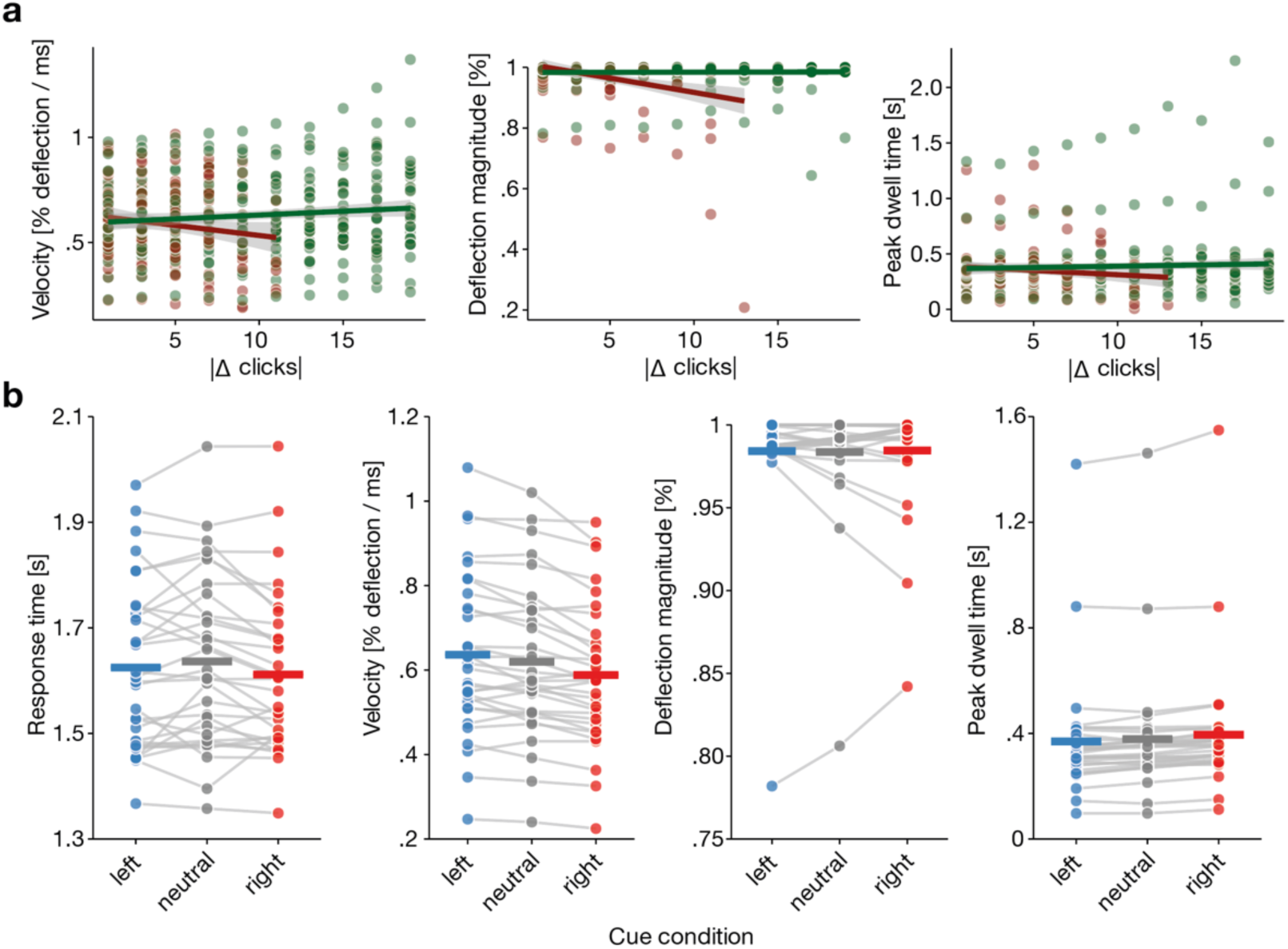
Descriptive overview of joystick metrics as a function of evidence strength in correct (green) and error (red) trials (**a**), and cue information (**b**). Individual coloured dots show participant-specific means, thick lines the group-level grand average.

**Figure S2:**
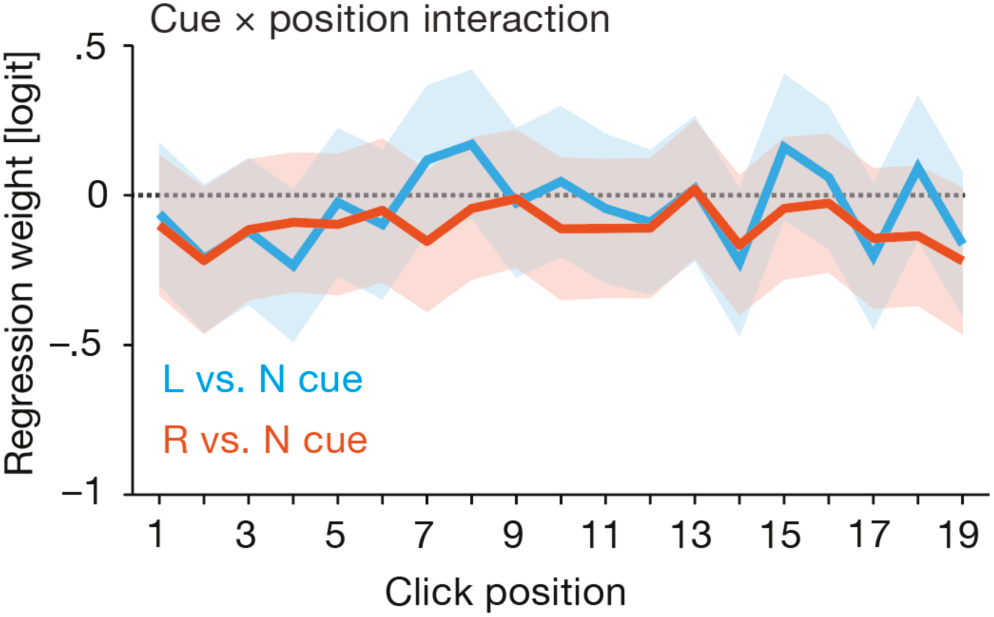
Cue x click position interaction kernels. Error bands indicate the 95 % confidence interval around logistic mixed-effect model derived regression weights per click position for the contrast of left vs. neutral cue (light blue) and right vs. neutral cue (orange).

**Figure S3:**
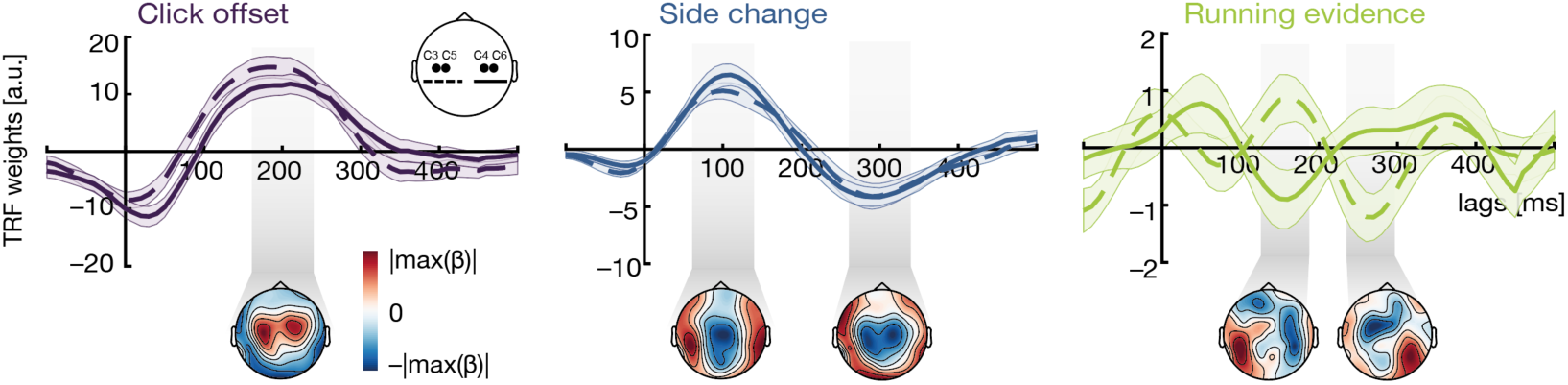
Grand average temporal response functions (TRFs) for regressors capturing final click offset, perceptual (side) change, and running evidence accumulation.

**Figure S4:**
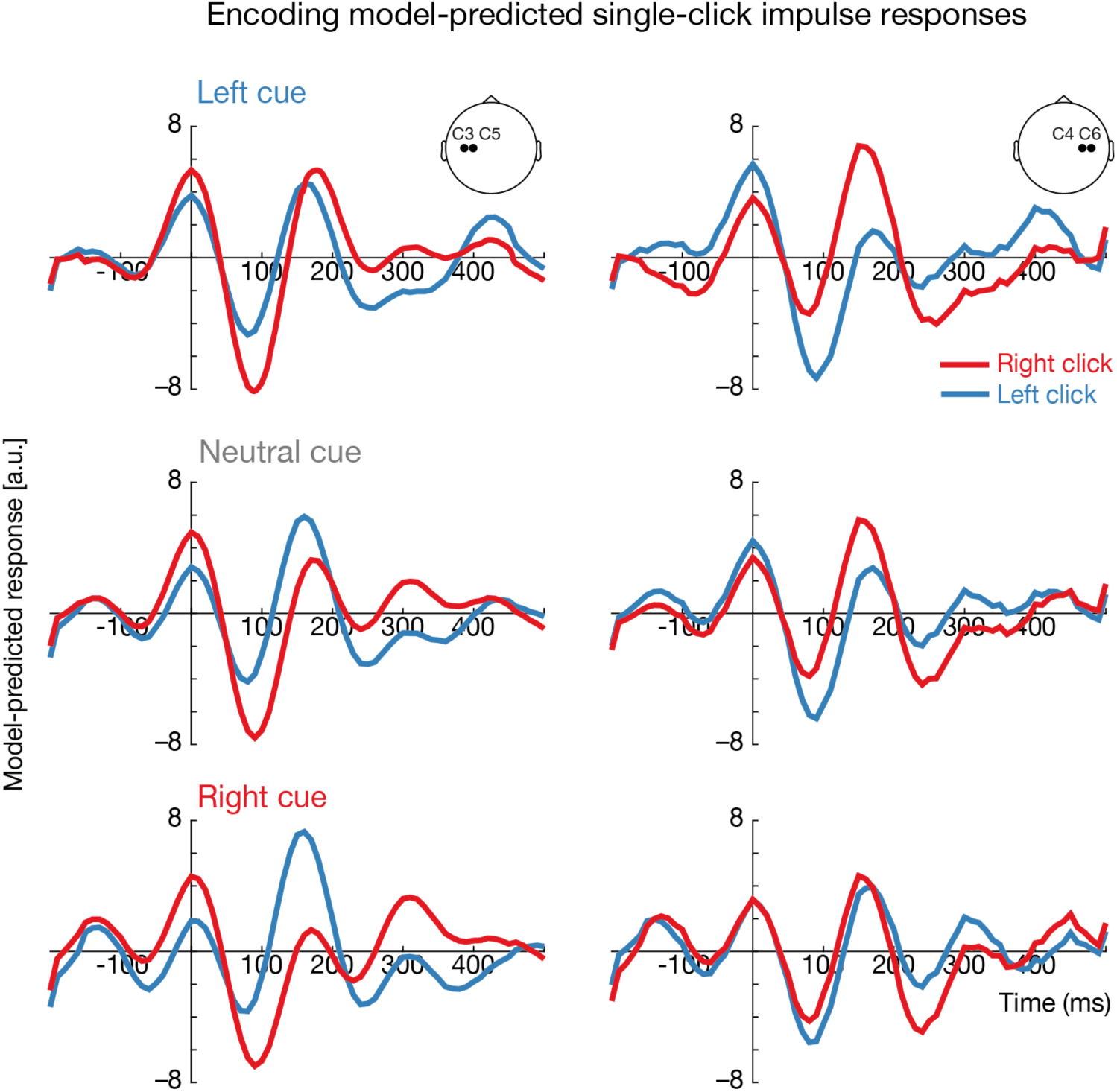
Neural encoding model-predicted impulse responses to a single click at lateralized auditory channels for different combinations of presentation side and cue condition.

**Figure S5:**
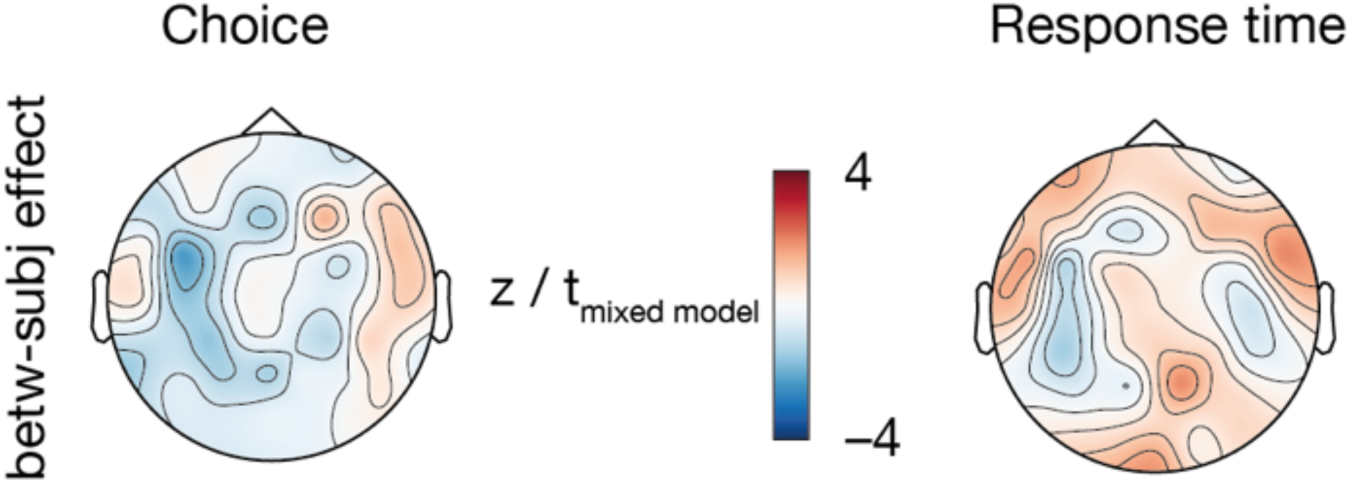
Between-subject effect of mean encoding strength on choice (left) and response time (right).

### Supplemental tables

**Table S1:**
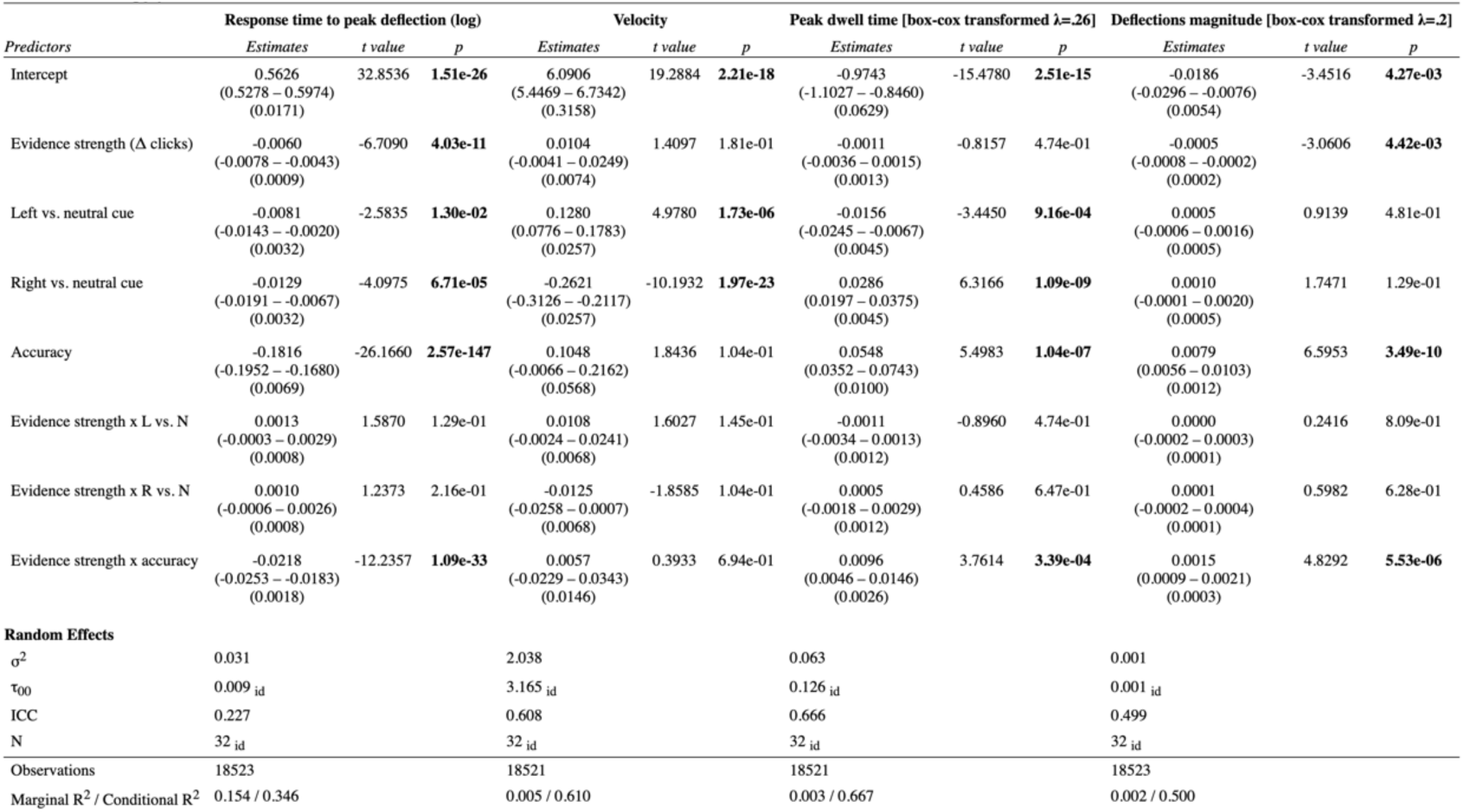
Modelling joystrick movement-based behavioural metrics.

**Tables S2:**
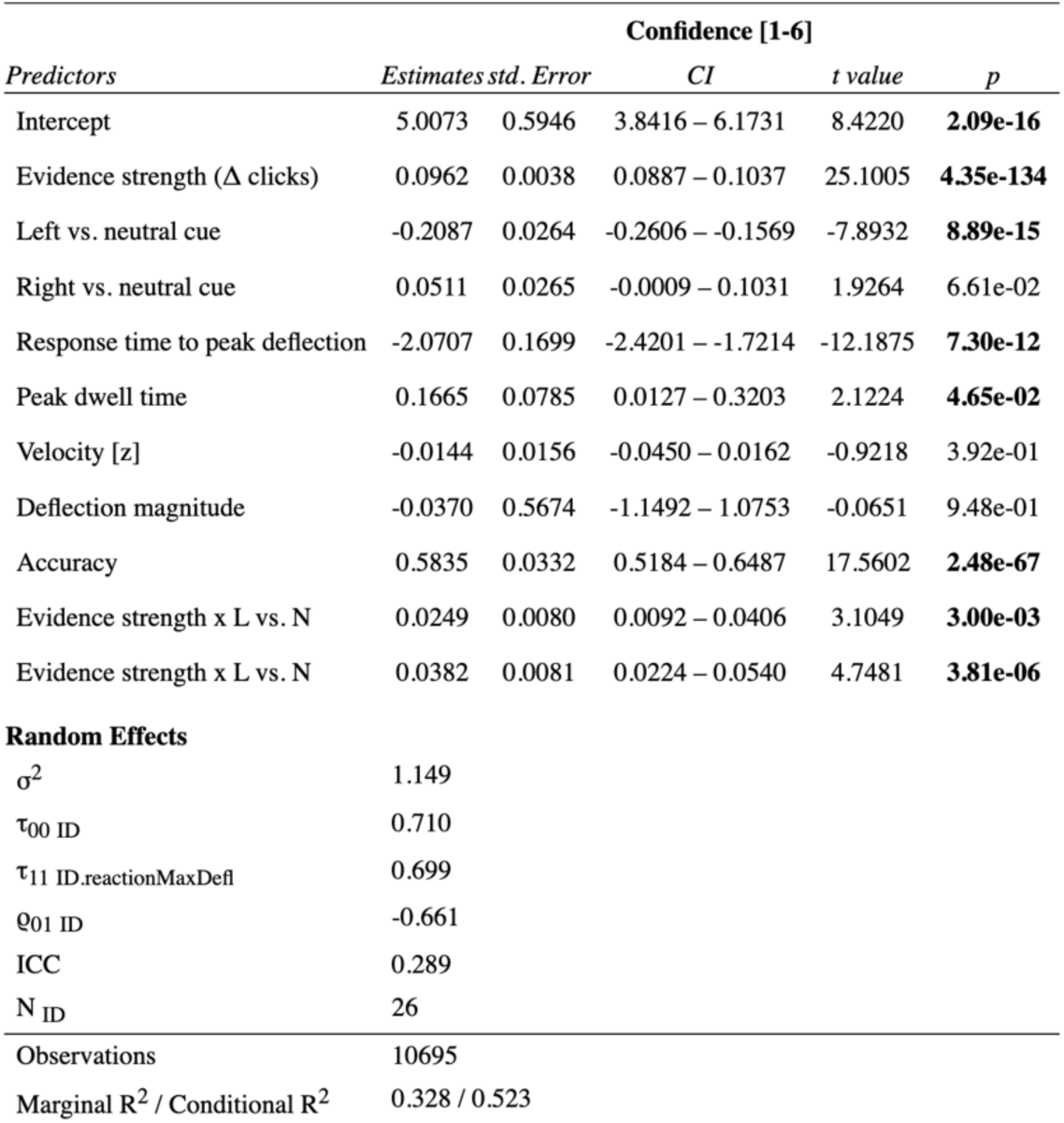
Modelling confidence.

**Table S3:** Modelling choice.

| Predictors | Odds Ratios | p('right') |  |  |
| --- | --- | --- | --- | --- |
|  |  | CI | z value | p |
| Intercept | 0.868 | 0.743 – 1.013 | -1.802 | 8.582e-02 |
| Evidence strength ($\Delta$ clicks) | 1.623 | 1.552 – 1.697 | 21.222 | **3.585e-99** |
| Left vs. neutral cue | 0.618 | 0.498 – 0.766 | -4.383 | **3.512e-05** |
| Right vs. neutral cue | 1.628 | 1.303 – 2.034 | 4.294 | **3.512e-05** |
| Evidence strength x L vs. N | 1.001 | 0.966 – 1.037 | 0.041 | 9.670e-01 |
| Evidence strength x R vs. N | 0.951 | 0.921 – 0.982 | -3.044 | **3.501e-03** |
| **Random Effects** |  |  |  |  |
| $\sigma^2$ | 3.290 | | | |
| $\tau_{00}$ id | 0.173 | | | |
| $\tau_{11}$ id.SNR | 0.014 | | | |
| $\tau_{11}$ id.Cues (left vs. neutr) | 0.249 | | | |
| $\tau_{11}$ id.Cues (right vs. neutr) | 0.287 | | | |
| Q01 | 0.001 |  |  |  |
|  | -0.013 |  |  |  |
|  | -0.226 |  |  |  |
| ICC | 0.227 |  |  |  |
| N id | 32 |  |  |  |
| Observations | 18523 |  |  |  |
| Marginal $R^2$ / Conditional $R^2$ | 0.747 / 0.804 | | | |

**Table S4:**
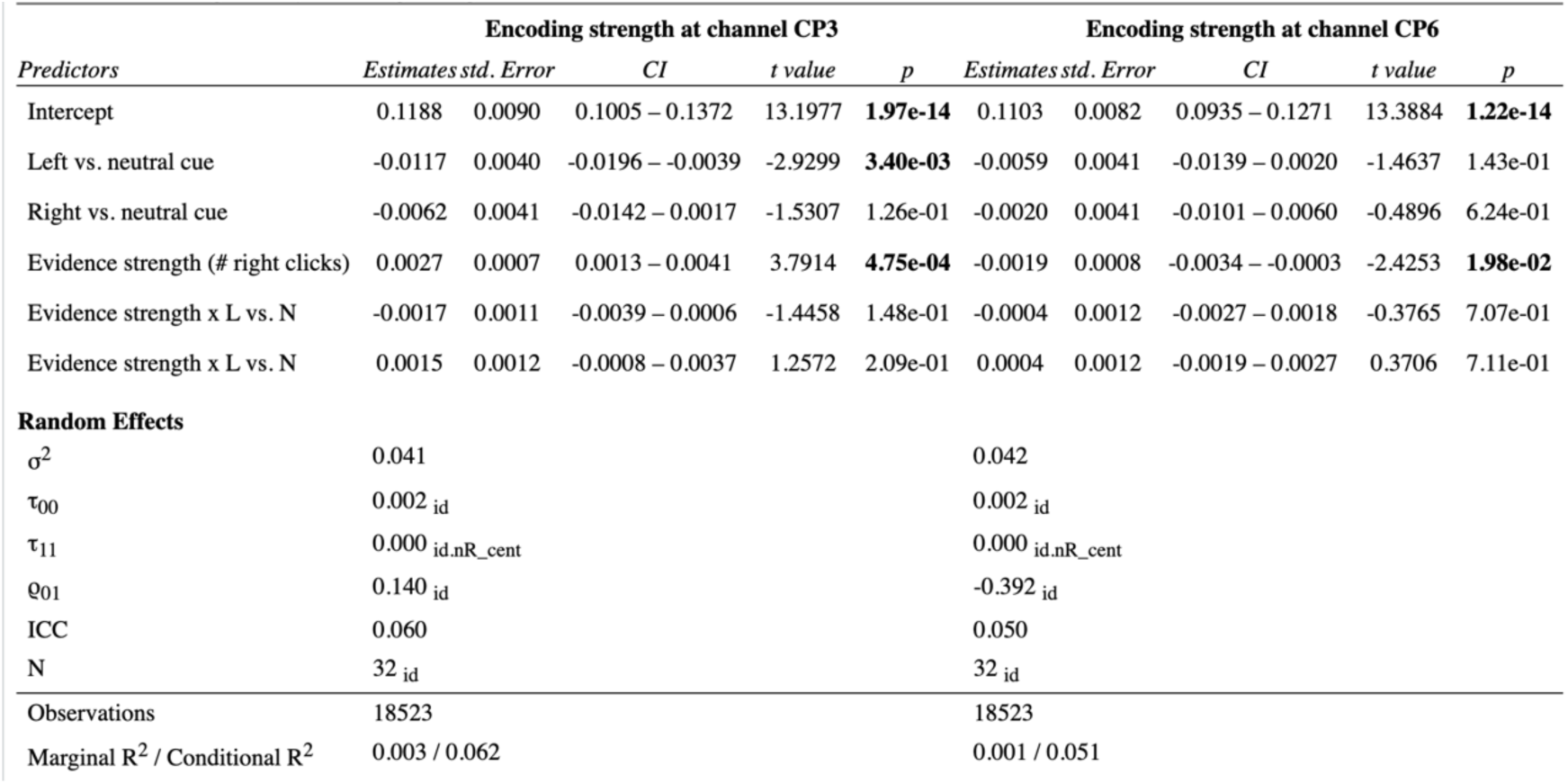
Modelling sensory encoding strength.

| Predictors | Encoding strength at channel CP3 |  |  |  |  | Encoding strength at channel CP6 |  |  |  |  |
| --- | --- | --- | --- | --- | --- | --- | --- | --- | --- | --- |
|  | Estimates | std. Error | CI | t value | p | Estimates | std. Error | CI | t value | p |
| Intercept | 0.1188 | 0.0090 | 0.1005 – 0.1372 | 13.1977 | **1.97e-14** | 0.1103 | 0.0082 | 0.0935 – 0.1271 | 13.3884 | **1.22e-14** |
| Left vs. neutral cue | -0.0117 | 0.0040 | -0.0196 – -0.0039 | -2.9299 | **3.40e-03** | -0.0059 | 0.0041 | -0.0139 – 0.0020 | -1.4637 | 1.43e-01 |
| Right vs. neutral cue | -0.0062 | 0.0041 | -0.0142 – 0.0017 | -1.5307 | 1.26e-01 | -0.0020 | 0.0041 | -0.0101 – 0.0060 | -0.4896 | 6.24e-01 |
| Evidence strength (# right clicks) | 0.0027 | 0.0007 | 0.0013 – 0.0041 | 3.7914 | **4.75e-04** | -0.0019 | 0.0008 | -0.0034 – -0.0003 | -2.4253 | **1.98e-02** |
| Evidence strength x L vs. N | -0.0017 | 0.0011 | -0.0039 – 0.0006 | -1.4458 | 1.48e-01 | -0.0004 | 0.0012 | -0.0027 – 0.0018 | -0.3765 | 7.07e-01 |
| Evidence strength x R vs. N | 0.0015 | 0.0012 | -0.0008 – 0.0037 | 1.2572 | 2.09e-01 | 0.0004 | 0.0012 | -0.0019 – 0.0027 | 0.3706 | 7.11e-01 |
| **Random Effects** |  |  |  |  |  |  |  |  |  |  |
| $\sigma^2$ | 0.041 | | | | | 0.042 | | | | |
| $\tau_{00}$ | 0.002 id | | | | | 0.002 id | | | | |
| $\tau_{11}$ | 0.000 id.nR_cent | | | | | 0.000 id.nR_cent | | | | |
| Q01 | 0.140 id |  |  |  |  | -0.392 id |  |  |  |  |
| ICC | 0.060 |  |  |  |  | 0.050 |  |  |  |  |
| N | 32 id |  |  |  |  | 32 id |  |  |  |  |
| Observations | 18523 |  |  |  |  | 18523 |  |  |  |  |
| Marginal $R^2$ / Conditional $R^2$ | 0.003 / 0.062 | | | | | 0.001 / 0.051 | | | | |

**Table S5:** Modelling choice from neural dynamics.

| Predictors | Odds Ratios | p('right') |  |  |
| --- | --- | --- | --- | --- |
|  |  | CI | z value | p |
| Intercept | 0.880 | 0.755 – 1.025 | -1.645 | 1.57e-01 |
| Evidence strength ($\Delta$ clicks) | 1.562 | 1.539 – 1.585 | 59.713 | **0.00e+00** |
| Left vs. neutral cue | 0.621 | 0.549 – 0.702 | -7.598 | **1.11e-13** |
| Right vs. neutral cue | 1.673 | 1.486 – 1.884 | 8.512 | **9.41e-17** |
| Neural encoding strength [within-subj] | 1.750 | 1.346 – 2.275 | 4.177 | **8.13e-05** |
| Neural encoding strength [between-subj] | 0.087 | 0.004 – 1.819 | -1.574 | 1.59e-01 |
| Parietal pre-stimulus ALI [within-subj] | 0.999 | 0.864 – 1.155 | -0.017 | 9.86e-01 |
| Parietal pre-stimulus ALI [between-subj] | 0.973 | 0.928 – 1.020 | -1.135 | 3.13e-01 |
| Evidence strength x L vs. N | 0.991 | 0.957 – 1.026 | -0.522 | 6.62e-01 |
| Evidence strength x R vs. N | 0.942 | 0.913 – 0.973 | -3.678 | **5.17e-04** |
| Neural encoding strength x ALI [both within-subj] | 1.254 | 0.994 – 1.582 | 1.908 | 1.03e-01 |
| **Random Effects** |  |  |  |  |
| $\sigma^2$ | 3.290 | | | |
| $\tau_{00}$ id | 0.171 | | | |
| $\tau_{11}$ id.Ch_CP3_within | 0.118 | | | |
| $\theta_{01}$ id | 0.187 | | | |
| ICC | 0.051 |  |  |  |
| N<sub>id</sub> | 32 |  |  |  |
| Observations | 18523 |  |  |  |
| Marginal R<sup>2</sup> / Conditional R<sup>2</sup> | 0.758 / 0.771 |  |  |  |

**Table S6:**
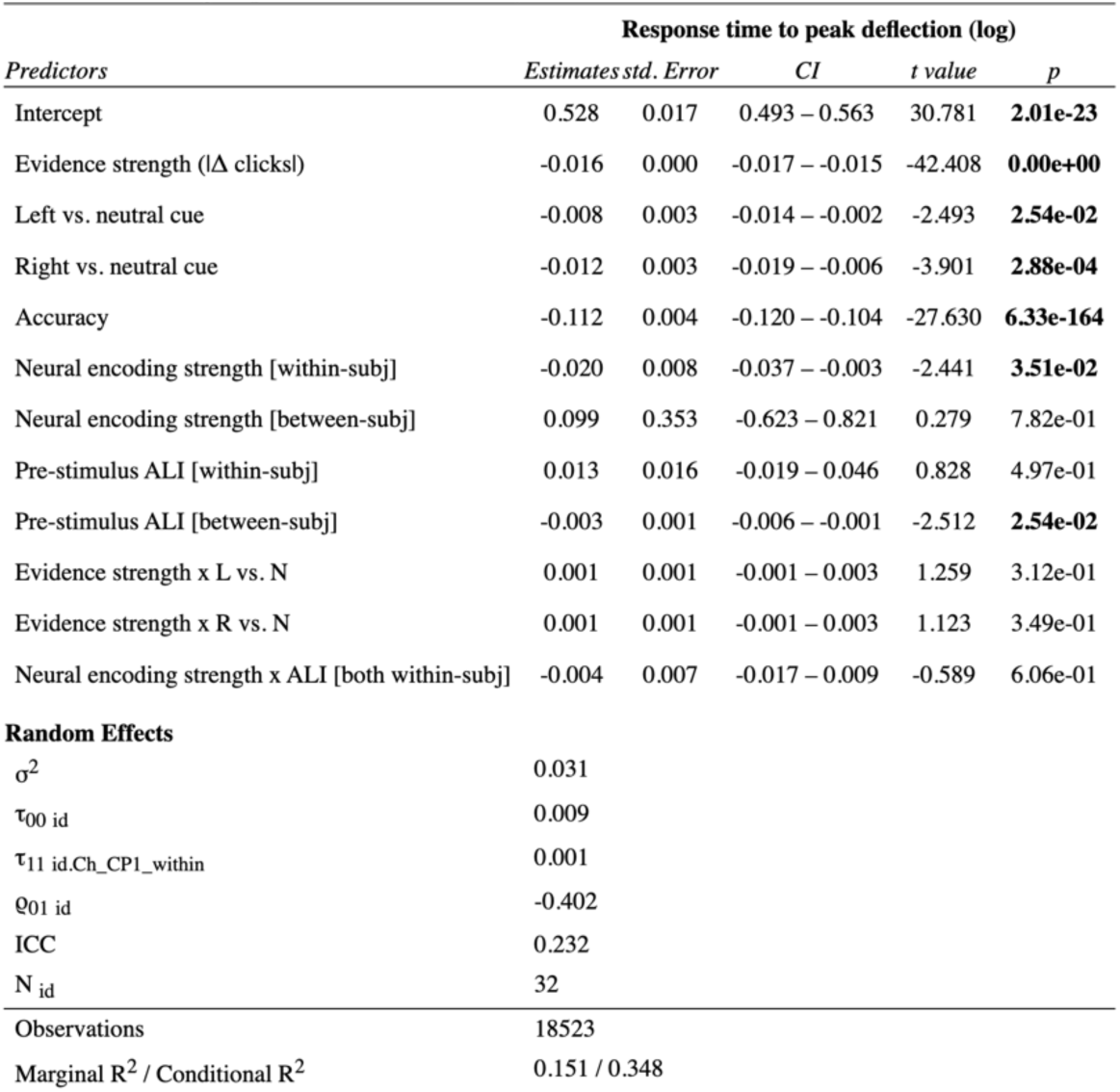
Modelling response time from neural dynamics.

## References

Adams, R. A., Huys, Q. J., & Roiser, J. P. (2016). Computational Psychiatry: towards a mathematically informed understanding of mental illness. J Neurol Neurosurg Psychiatry, 87(1), 53–63. 10.1136/jnnp-2015-310737

Adams, R. A., Napier, G., Roiser, J. P., Mathys, C., & Gilleen, J. (2018). Attractor-like Dynamics in Belief Updating in Schizophrenia. J Neurosci, 38(44), 9471– 9485. 10.1523/JNEUROSCI.3163-17.2018

Alavash, M., Tune, S., & Obleser, J. (2019). Modular reconfiguration of an auditory control brain network supports adaptive listening behavior. Proceedings of the National Academy of Science of the United States of America, 116(2), 660–669. 10.1073/pnas.1815321116

Bell, A., Fairbrother, M., & Jones, K. (2018). Fixed and random effects models: making an informed choice [OriginalPaper]. Quality & Quantity, 53(2), 1051–1074. 10.1007/s11135-018-0802-x

Bogacz, R., Brown, E., Moehlis, J., Holmes, P., & Cohen, J. D. (2006). The physics of optimal decision making: A formal analysis of models of performance in two-alternative forced-choice tasks. Psychological review, 113(4), 700–765. 10.1037/0033-295x.113.4.700

Brown, S., & Nicholls, M. E. R. (1997). Hemispheric asymmetries for the temporal resolution of brief auditory stimuli. Perception & Psychophysics, 59(3), 442–447. 10.3758/BF03211910

Brunton, B. W., Botvinick, M. M., & Brody, C. D. (2013). Rats and humans can optimally accumulate evidence for decision-making. Science, 340(6128), 95–98. 10.1126/science.1233912

Buxbaum, L. J., Kyle, K., Grossman, M., & Coslett, B. (2007). Left Inferior Parietal Representations for Skilled Hand-Object Interactions: Evidence from Stroke and Corticobasal Degeneration. CORTEX, 43(3), 411–423. 10.1016/S0010-9452(08)70466-0

Carandini, M., & Heeger, D. J. (2011). Normalization as a canonical neural computation. [Review]. Nature Reviews Neuroscience, 13(1), 51–62. 10.1038/nrn3136

Cheadle, S., Wyart, V., Tsetsos, K., Myers, N., de Gardelle, V., Herce Castanon, S., & Summerfield, C. (2014). Adaptive gain control during human perceptual choice. Neuron, 81(6), 1429–1441. 10.1016/j.neuron.2014.01.020

Cortes-Briones, J., Urrutia-Gandolfo, J., Estevez, P. A., Sengupta, A., Basilico, M., Skosnik, P., Limoncelli, D., D’Souza, D. C., Petrakis, I., & Krystal, J. (2025). Opposing Modulation of EEG Aperiodic Component by Ketamine and Thiopental: Implications for the Noninvasive Assessment of Cortical E/I Balance in Humans. bioRxiv. 10.1101/2025.05.12.653580

Crosse, M. J., Di Liberto, G. M., Bednar, A., & Lalor, E. C. (2016). The Multivariate Temporal Response Function (mTRF) Toolbox: A MATLAB Toolbox for Relating Neural Signals to Continuous Stimuli. Front Hum Neurosci, 10, 604. 10.3389/fnhum.2016.00604

Crosse, M. J., Zuk, N. J., Di Liberto, G. M., Nidiffer, A. R., Molholm, S., & Lalor, E. C. (2021). Linear Modeling of Neurophysiological Responses to Speech and Other Continuous Stimuli: Methodological Considerations for Applied Research. Front Neurosci, 15, 705621. 10.3389/fnins.2021.705621

Crowley, K. E., & Colrain, I. M. (2004). A review of the evidence for P2 being an independent component process: Age, sleep and modality. Clinical Neurophysiology, 115(4), 732–744. 10.1016/j.clinph.2003.11.021

de Gardelle, V., & Summerfield, C. (2011). Robust averaging during perceptual judgment. Proc Natl Acad Sci U S A, 108(32), 13341–13346. 10.1073/pnas.1104517108

de Lange, F. P., Jensen, O., & Dehaene, S. (2010). Accumulation of evidence during sequential decision making: the importance of top-down factors. J Neurosci, 30(2), 731–738. 10.1523/JNEUROSCI.4080-09.2010

de Lange, F. P., Rahnev, D. A., Donner, T. H., & Lau, H. (2013). Prestimulus Oscillatory Activity over Motor Cortex Reflects Perceptual Expectations. Journal of Neuroscience, 33(4), 1400–1410. 10.1523/JNEUROSCI.1094-12.2013

Donner, T. H., Siegel, M., Fries, P., & Engel, A. K. (2009). Buildup of choice-predictive activity in human motor cortex during perceptual decision making. Curr Biol, 19(18), 1581–1585. 10.1016/j.cub.2009.07.066

Egner, T., Monti, J. M., & Summerfield, C. (2010). Expectation and Surprise Determine Neural Population Responses in the Ventral Visual Stream. The Journal of Neuroscience.

Erb, J., Kreitewolf, J., Pinheiro, A., & Obleser, J. (2020). Aberrant perceptual judgments on speech-relevant acoustic features in hallucination-prone individuals. 1–15. 10.1093/schizbullopen/sgaa059

Erb, J., & Obleser, J. (2020). Neural filters for challening listening situations. In M. Gazzaniga, G. R. Mangun, & D. Poeppel (Eds.), The cognitive neurosciences (6th ed.). MIT Press.

Ernst, M. O., & Banks, M. S. (2002). Humans integrate visual and haptic information in a statistically optimal fashion. Nature, 415(6870), 429–433. 10.1038/415429a

Forstmann, B. U., Anwander, A., Schafer, A., Neumann, J., Brown, S., Wagenmakers, E. J., Bogacz, R., & Turner, R. (2010). Cortico-striatal connections predict control over speed and accuracy in perceptual decision making. Proc Natl Acad Sci U S A, 107(36), 15916–15920. 10.1073/pnas.1004932107

Foxe, J. J., & Snyder, A. C. (2011). The Role of Alpha-Band Brain Oscillations as a Sensory Suppression Mechanism during Selective Attention. Frontiers in Psychology, 2, 154. 10.3389/fpsyg.2011.00154

Garrett, D. D., Kovacevic, N., McIntosh, A. R., & Grady, C. L. (2011). The Importance of Being Variable. Journal of Neuroscience, 31(12), 4496–4503. 10.1523/JNEUROSCI.5641-10.2011

Garrett, D. D., Samanez-Larkin, G. R., MacDonald, S. W., Lindenberger, U., McIntosh, A. R., & Grady, C. L. (2013). Moment-to-moment brain signal variability: a next frontier in human brain mapping? Neurosci Biobehav Rev, 37(4), 610–624. 10.1016/j.neubiorev.2013.02.015

Gold, J. I., & Shadlen, M. N. (2007). The neural basis of decision making. Annu Rev Neurosci, 30, 535–574. 10.1146/annurev.neuro.29.051605.113038

Gould, I. C., Nobre, A. C., Wyart, V., & Rushworth, M. F. (2012). Effects of decision variables and intraparietal stimulation on sensorimotor oscillatory activity in the human brain. J Neurosci, 32(40), 13805–13818. 10.1523/JNEUROSCI.2200-12.2012

Gupta, D., DePasquale, B., Kopec, C. D., & Brody, C. D. (2024). Trial-history biases in evidence accumulation can give rise to apparent lapses in decision-making. Nat Commun, 15(1), 662. 10.1038/s41467-024-44880-5

Hanks, T. D., Kopec, C. D., Brunton, B. W., Duan, C. A., Erlich, J. C., & Brody, C. D. (2015). Distinct relationships of parietal and prefrontal cortices to evidence accumulation. Nature, 520(7546), 220–223. 10.1038/nature14066

Henry, M. J., & Obleser, J. (2012). Frequency modulation entrains slow neural oscillations and optimizes human listening behavior. Proceedings of the National Academy of Science of the United States of America, 109(49), 20095–20100. 10.1073/pnas.1213390109

Iemi, L., Chaumon, M., Crouzet, S. M., & Busch, N. A. (2017). Spontaneous Neural Oscillations Bias Perception by Modulating Baseline Excitability. J Neurosci, 37(4), 807–819. 10.1523/JNEUROSCI.1432-16.2016

Iemi, L., Gwilliams, L., Samaha, J., Auksztulewicz, R., Cycowicz, Y. M., King, J.-R., Nikulin, V. V., Thesen, T., Doyle, W., Devinsky, O., Schroeder, C. E., Melloni, L., & Haegens, S. (2021). Spontaneous neural oscillations influence behavior and sensory representations by suppressing neuronal excitability. Nature Communications, 2021.2003.2001.433450. 10.1101/2021.03.01.433450

Jardri, R., & Deneve, S. (2013). Circular inferences in schizophrenia. Brain, 136(Pt 11), 3227–3241. 10.1093/brain/awt257

Kayser, J., & Tenke, C. E. (2015). Issues and considerations for using the scalp surface Laplacian in EEG/ERP research: A tutorial review. Int J Psychophysiol, 97(3), 189–209. 10.1016/j.ijpsycho.2015.04.012

Kerlin, J. R., Shahin, A. J., & Miller, L. M. (2010). Attentional gain control of ongoing cortical speech representations in a "cocktail party". The Journal of Neuroscience, 30(2), 620–628. 10.1523/JNEUROSCI.3631-09.2010

Keung, W., Hagen, T. A., & Wilson, R. C. (2019). Regulation of evidence accumulation by pupil-linked arousal processes. Nature Human Behaviour, 1–12. 10.1038/s41562-019-0551-4

Keung, W., Hagen, T. A., & Wilson, R. C. (2020). A divisive model of evidence accumulation explains uneven weighting of evidence over time. Nat Commun, 11(1), 2160. 10.1038/s41467-020-15630-0

Kok, P., Brouwer, G. J., van Gerven, M. A., & de Lange, F. P. (2013). Prior expectations bias sensory representations in visual cortex. J Neurosci, 33(41), 16275– 16284. 10.1523/JNEUROSCI.0742-13.2013

Krakauer, J. W., Ghazanfar, A. A., Gomez-Marin, A., MacIver, M. A., & Poeppel, D. (2017). Neuroscience needs behavior: correcting a reductionist bias. Neuron, 93(3), 480–490. 10.1016/j.neuron.2016.12.041

Kunze, J., Student, J., Tune, S., Herrmann, A., Göttlich, M., Oster, H., Tzabazis, A., Lorenz, B., Nau, C., & Obleser, J. (2025). Low doses of ketamine and propofol shift multimodal noninvasive metrics of excitation-inhibition balance and cardiovascular response. Society for Neuroscience Meeting, San Diego.

Lalor, E. C., Power, A. J., Reilly, R. B., & Foxe, J. J. (2009). Resolving precise temporal processing properties of the auditory system using continuous stimuli. J Neurophysiol, 102(1), 349–359. 10.1152/jn.90896.2008

Lam, N. H., Borduqui, T., Hallak, J., Roque, A., Anticevic, A., Krystal, J. H., Wang, X. J., & Murray, J. D. (2022). Effects of Altered Excitation-Inhibition Balance on Decision Making in a Cortical Circuit Model. J Neurosci, 42(6), 1035–1053. 10.1523/JNEUROSCI.1371-20.2021

Linares, D., & López-Moliner, J. (2016). quickpsy: An R Package to Fit Psychometric Functions for Multiple Groups. The R Journal, 8, 122–131.

Luck, S. J., Hillyard, S. A., Mouloua, M., & Hawkins, H. L. (1996). Mechanisms of visual–spatial attention: Resource allocation or uncertainty reduction? Journal of Experimental Psychology: Human Perception and Performance, 22(3), 725–737. 10.1037/0096-1523.22.3.725

Luke, S. G. (2017). Evaluating significance in linear mixed-effects models in R. Behavior Research Methods, 49(4), 1494–1502. 10.3758/s13428-016-0809-y

Mulder, M. J., Wagenmakers, E. J., Ratcliff, R., Boekel, W., & Forstmann, B. U. (2012). Bias in the brain: a diffusion model analysis of prior probability and potential payoff. J Neurosci, 32(7), 2335–2343. 10.1523/JNEUROSCI.4156-11.2012

Müller, N., & Weisz, N. (2011). Lateralized auditory cortical alpha band activity and interregional connectivity pattern reflect anticipation of target sounds. Cerebral Cortex, 22(7), 1604–1613. 10.1093/cercor/bhr232

Murphy, P. R., Wilming, N., Hernandez-Bocanegra, D. C., Prat-Ortega, G., & Donner, T. H. (2021). Adaptive circuit dynamics across human cortex during evidence accumulation in changing environments. Nat Neurosci, 24(7), 987–997. 10.1038/s41593-021-00839-z

Murray, J. D., & Wang, X.-J. (2018). Cortical Circuit Models in Psychiatry. In Computational Psychiatry (pp. 3–25). 10.1016/b978-0-12-809825-7.00001-8

Musall, S., Kaufman, M. T., Juavinett, A. L., Gluf, S., & Churchland, A. K. (2019). Single-trial neural dynamics are dominated by richly varied movements. Nature Neuroscience, 22(10), 1–16. 10.1038/s41593-019-0502-4

Musall, S., Sun, X. R., Mohan, H., An, X., Gluf, S., Li, S. J., Drewes, R., Cravo, E., Lenzi, I., Yin, C., Kampa, B. M., & Churchland, A. K. (2023). Pyramidal cell types drive functionally distinct cortical activity patterns during decision-making. Nat Neurosci, 26(3), 495–505. 10.1038/s41593-022-01245-9

Näätänen, R., Paavilainen, P., Rinne, T., & Alho, K. (2007). The mismatch negativity (MMN) in basic research of central auditory processing: A review. Clinical Neurophysiology, 118(12), 2544–2590. 10.1016/j.clinph.2007.04.026

Näätänen, R., & Picton, T. (1987). The N1 Wave of the Human Electric and Magnetic Response to Sound: A Review and an Analysis of the Component Structure. Psychophysiology, 24(4), 375–425. 10.1111/j.1469-8986.1987.tb00311.x

Obleser, J. (2025). Metacognition in the listening brain. Trends Neurosci, 48(2), 100–112. 10.1016/j.tins.2024.12.007

Okazawa, G., & Kiani, R. (2023). Neural Mechanisms That Make Perceptual Decisions Flexible. Annu Rev Physiol, 85, 191–215. 10.1146/annurev-physiol-031722-024731

Okazawa, G., Sha, L., Purcell, B. A., & Kiani, R. (2018). Psychophysical reverse correlation reflects both sensory and decision-making processes. [OriginalPaper]. Nature Communications, 9(1), 3479–3416. 10.1038/s41467-018-05797-y

Pagan, M., Tang, V. D., Aoi, M. C., Pillow, J. W., Mante, V., Sussillo, D., & Brody, C. D. (2024). Individual variability of neural computations underlying flexible decisions. Nature. 10.1038/s41586-024-08433-6

Payne, L., Rogers, C. S., Wingfield, A., & Sekuler, R. (2016). A right-ear bias of auditory selective attention is evident in alpha oscillations. Psychophysiology, 54(4), 528–535. 10.1111/psyp.12815

Pernet, C. R., Sajda, P., & Rousselet, G. A. (2011). Single-trial analyses: why bother? Frontiers in Psychology, 2, 322. 10.3389/fpsyg.2011.00322

Picton, T. (2013). Hearing in time: evoked potential studies of temporal processing. Ear Hear, 34(4), 385–401. 10.1097/AUD.0b013e31827ada02

Pion-Tonachini, L., Kreutz-Delgado, K., & Makeig, S. (2019). ICLabel: An automated electroencephalographic independent component classifier, dataset, and website. NeuroImage, 198, 181–197. 10.1016/j.neuroimage.2019.05.026

Plomp, R. (1964). Rate of Decay of Auditory Sensation. The Journal of the Acoustical Society of America, 36(2), 277–282. 10.1121/1.1918946

Powers, A. R., Mathys, C., & Corlett, P. R. (2017). Pavlovian conditioning–induced hallucinations result from overweighting of perceptual priors. Science, 357(6351), 596–600. 10.1126/science.aan3458

Rahnev, D., Lau, H., & de Lange, F. P. (2011). Prior expectation modulates the interaction between sensory and prefrontal regions in the human brain. Journal of Neuroscience, 31(29), 10741–10748. 10.1523/JNEUROSCI.1478-11.2011

Ratcliff, R., & McKoon, G. (2008). The diffusion decision model: theory and data for two-choice decision tasks. Neural Comput, 20(4), 873–922. 10.1162/neco.2008.12-06-420

Ratcliff, R., & Smith, P. L. (2004). A comparison of sequential sampling models for two-choice reaction time. Psychol Rev, 111(2), 333–367. 10.1037/0033-295X.111.2.333

Rockstroh, B., Müller, M., Cohen, R., & Elbet, T. (1992). Probing the functional brain state during P300-evocation. Journal of Psychophysiology, 6(2), 175–184.

Ross, B., & Tremblay, K. (2009). Stimulus experience modifies auditory neuromagnetic responses in young and older listeners. Hear Res, 248(1-2), 48–59. 10.1016/j.heares.2008.11.012

Salvador, A., Arnal, L. H., Vinckier, F., Domenech, P., Gaillard, R., & Wyart, V. (2022). Premature commitment to uncertain decisions during human NMDA receptor hypofunction. [OriginalPaper]. Nature Communications, 13(1), 338–315. 10.1038/s41467-021-27876-3

Samaha, J., Iemi, L., Haegens, S., & Busch, N. A. (2020). Spontaneous Brain Oscillations and Perceptual Decision-Making. Trends in Cognitive Sciences, 1–15. 10.1016/j.tics.2020.05.004

Sanders, J. I., Hangya, B., & Kepecs, A. (2016). Signatures of a Statistical Computation in the Human Sense of Confidence. Neuron, 90(3), 499–506. 10.1016/j.neuron.2016.03.025

Schroger, E., Marzecova, A., & SanMiguel, I. (2015). Attention and prediction in human audition: a lesson from cognitive psychophysiology. Eur J Neurosci, 41(5), 641–664. 10.1111/ejn.12816

Smith, P. L., & Ratcliff, R. (2004). Psychology and neurobiology of simple decisions. Trends Neurosci, 27(3), 161–168. 10.1016/j.tins.2004.01.006

Spitzer, B., Waschke, L., & Summerfield, C. (2017). Selective overweighting of larger magnitudes during noisy numerical comparison. Nat Hum Behav, 1(8), 145. 10.1038/s41562-017-0145

St John-Saaltink, E., Kok, P., Lau, H. C., & de Lange, F. P. (2016). Serial Dependence in Perceptual Decisions Is Reflected in Activity Patterns in Primary Visual Cortex. J Neurosci, 36(23), 6186–6192. 10.1523/JNEUROSCI.4390-15.2016

Stocker, A. A., & Simoncelli, E. P. (2006). Noise characteristics and prior expectations in human visual speed perception. Nat Neurosci, 9(4), 578–585. 10.1038/nn1669

Strauß, A., Wöstmann, M., & Obleser, J. (2014). Cortical alpha oscillations as a tool for auditory selective inhibition. Frontiers in human neuroscience, 8, 1–7. 10.3389/fnhum.2014.00350

Summerfield, C., & de Lange, F. P. (2014). Expectation in perceptual decision making: neural and computational mechanisms. Nature Publishing Group, 1–12. 10.1038/nrn3838

Summerfield, C., & Koechlin, E. (2008). A Neural Representation of Prior Information during Perceptual Inference. Neuron, 59(2), 336–347. 10.1016/j.neuron.2008.05.021

Szul, M. J., Bompas, A., Sumner, P., & Zhang, J. (2020). The validity and consistency of continuous joystick response in perceptual decision-making. Behav Res Methods, 52(2), 681–693. 10.3758/s13428-019-01269-3

Tanaka, K., Ross, B., Kuriki, S., Harashima, T., Obuchi, C., & Okamoto, H. (2021). Neurophysiological Evaluation of Right-Ear Advantage During Dichotic Listening. Front Psychol, 12, 696263. 10.3389/fpsyg.2021.696263

Tremblay, K. L., Ross, B., Inoue, K., McClannahan, K., & Collet, G. (2014). Is the auditory evoked P2 response a biomarker of learning? Front Syst Neurosci, 8, 28. 10.3389/fnsys.2014.00028

Tune, S., Alavash, M., Fiedler, L., & Obleser, J. (2021). Neural attentional-filter mechanisms of listening success in middle-aged and older individuals. Nature Communications, 1–14. 10.1038/s41467-021-24771-9

Tune, S., Kunze, J., Student, J., Nau, C., & Obleser, J. (2025). Perturbing the neurophysiological excitation-inhibition balance of auditory decision-making Society for Neuroscience Meeting, San Diego.

Tune, S., & Obleser, J. (2024). Neural attentional filters and behavioural outcome follow independent individual trajectories over the adult lifespan. eLife, 12. 10.7554/eLife.92079

Tune, S., Wöstmann, M., & Obleser, J. (2018). Probing the limits of alpha power lateralisation as a neural marker of selective attention in middle-aged and older listeners. European Journal of Neuroscience, 48(7), 2537–2550. 10.1111/ejn.13862

von Helmholtz, H. (1867). Handbuch der physiologischen Optik.

Walsh, K., McGovern, D. P., Dully, J., Kelly, S. P., & O’Connell, R. G. (2024). Prior probability cues bias sensory encoding with increasing task exposure. eLife, 12. 10.7554/eLife.91135

Waschke, L., Kloosterman, N. A., Obleser, J., & Garrett, D. D. (2021). Behavior needs neural variability. [Review]. Neuron, 109(5), 751–766. 10.1016/j.neuron.2021.01.023

Waschke, L., Tune, S., & Obleser, J. (2019). Local cortical desynchronization and pupil-linked arousal differentially shape brain states for optimal sensory performance. eLife, 8, 1868–1827. 10.7554/eLife.51501

Waschke, L., Wöstmann, M., & Obleser, J. (2017). States and traits of neural irregularity in the age-varying human brain. 7(1), 1–12. 10.1038/s41598-017-17766-4

Waskom, M. L., Okazawa, G., & Kiani, R. (2019). Designing and Interpreting Psychophysical Investigations of Cognition. 1–13. 10.1016/j.neuron.2019.09.016

Weiss, Y., Simoncelli, E. P., & Adelson, E. H. (2002). Motion illusions as optimal percepts. Nature Neuroscience, 5(6), 598–604. 10.1038/nn0602-858

Wimmer, K., Compte, A., Roxin, A., Peixoto, D., Renart, A., & de la Rocha, J. (2015). Sensory integration dynamics in a hierarchical network explains choice probabilities in cortical area MT. Nat Commun, 6, 6177. 10.1038/ncomms7177

Woldorff, M., Hansen, J. C., & Hillyard, S. A. (1987). Evidence for effects of selective attention in the mid-latency range of the human auditory event-related potential. Electroencephalogr Clin Neurophysiol Suppl, 40, 146–154.

Wöstmann, M., Herrmann, B., Maess, B., & Obleser, J. (2016). Spatiotemporal dynamics of auditory attention synchronize with speech. Proceedings of the National Academy of Sciences, 113(14), 3873–3878. 10.1073/pnas.1523357113

Wöstmann, M., Waschke, L., & Obleser, J. (2019). Prestimulus neural alpha power predicts confidence in discriminating identical auditory stimuli. European Journal of Neuroscience, 49(1), 94–105. 10.1111/ejn.14226

Wu, M. C. K., David, S. V., & Gallant, J. L. (2006). Complete functional characterization of sensory neurons by system identification (Vol. 29) [Review]. 10.1146/annurev.neuro.29.051605.113024

Wyart, V., de Gardelle, V., Scholl, J., & Summerfield, C. (2012). Rhythmic Fluctuations in Evidence Accumulation during Decision Making in the Human Brain. Neuron, 76(4), 847–858. 10.1016/j.neuron.2012.09.015

Wyart, V., Nobre, A. C., & Summerfield, C. (2012). Dissociable prior influences of signal probability and relevance on visual contrast sensitivity. Proceedings of the National Academy of Sciences of the United States of America, 109(9), 3593–3598. 10.2307/41506995?ref=search-gateway:c65adb21b8f6eeb97e90e68b5d3e833a

